# Multispecific nanobody degraders co-deplete membrane receptors and enable targeted delivery of diverse payloads

**DOI:** 10.64898/2026.05.02.722401

**Authors:** Md Kabir, Yong Joon (Jeffrey) Kim, Zhijie Deng, Yufei Xiang, Paul R. Sargunas, Nuozi Song, Zishan Wang, Nesteene J. Param, Changzhong Jin, Zhe Sang, Alicia Yue, Alba Bundo, Razeen Hossain, Yue Zhong, Yindan Lin, Yan Xiong, Ernesto Guccione, Kuan-lin Huang, Mingye Feng, Jian Jin, Yi Shi

## Abstract

Targeting membrane receptors underlies the success of antibody-drug conjugates (ADCs), yet single-receptor formats can be limited by heterogeneous expression, compensatory signaling, and variable internalization. Here we developed Multivalent Interchangeable Nanobody Degradation System (MINDS), a modular nanobody-Fc chassis that co-engages multiple membrane receptors, promotes their lysosomal co-depletion, and enables delivery of diverse intracellular payloads. As a proof of concept, we generated Tritazumab, a trispecific nanobody-Fc targeting three oncogenic receptors EGFR, cMET, and TfR1. Tritazumab incorporates a high-affinity, non-transferrin-competing anti-TfR1 nanobody that drives efficient uptake and lysosomal trafficking, enabling coordinated depletion of all three receptors. Across non-small cell lung cancer models, Tritazumab achieved rapid and sustained multi-receptor surface loss with picomolar degradation potency, reaching near-maximal depletion within approximately 1.5 hours. Conjugation of Tritazumab to MMAE preserved receptor binding and produced substantially greater antiproliferative activity and improved tumor selectivity relative to clinical ADCs in matched cell models, along with potent *in viv*o tumor growth inhibition and acceptable tolerability in a xenograft model. Extending the platform beyond cytotoxic payloads, a BRD4 molecular glue conjugate improved the selectivity window by > 100-fold and showed marked *in vivo* efficacy, while an EZH2-targeting PROTAC conjugate achieved an approximately 1,000-fold increase in intracellular degradation potency relative to the free PROTAC. These findings establish MINDS as a modular multispecific degrader-payload platform that integrates receptor co-depletion to enhance anticancer selectivity and efficacy.

## Main

Antibody-drug conjugates (ADCs) have substantially advanced cancer therapy by coupling antibody targeting with potent small-molecule payloads, improving selectivity over systemically delivered cytotoxic chemotherapy and expanding the set of druggable antigens^1–3^. ADCs enter cells via receptor-mediated endocytosis, and payloads are released in the lysosomal pathway through pH- or enzyme-dependent linker cleavage^4^. In most ADCs, tumor cell killing is driven by the released payload, with the antibody primarily directing selective uptake. ADCs generally display higher anticancer efficacies than unconjugated monoclonal antibodies (mAbs) and improved safety relative to stand-alone chemotherapies^5^. As of 2024, 15 ADCs have been approved, with hundreds of trials underway globally ^6,7^.

Clinical performance nevertheless varies because mAb internalization efficiency is often constrained by the intrinsic trafficking properties of the target receptor together with antibody-specific features such as affinity and epitope.^3^ Many receptors and antibodies are therefore inefficient shuttles. Moreover, monospecific ADCs impose strong selective pressure on tumors, which can adapt via antigen downregulation, epitope masking, drug resistant mutations, or pathway rewiring, leading to outgrowth of resistant subclones ^8^. Together with heterogeneous antigen expression and compensatory signaling through parallel pathways, these features limit breadth and durability and can contribute to on-target, off-tumor toxicity when tumor-to-normal expression differentials are modest ^9^. Multispecific IgG-based formats may broaden coverage and improve uptake, but they often require complex multi-arm architectures that are difficult to generalize across targets and payloads and adds manufacturing complexity^10–12^.

In parallel, emerging extracellular degrader modalities, including lysosome-targeting chimeras/LYTACs, antibody-based PROTAC, cytokine receptor or folate receptor-targeting chimeras, have established receptor depletion as a viable alternative to receptor inhibition, with distinct kinetics and pharmacologic consequences^13–20^. However, most current implementations act on a single receptor, often with limited degradation efficacy, and/or rely on routing pathways that are not tumor-preferential, therefore restricting coordinated target control and translatability.

Here we introduce the Multivalent Interchangeable Nanobody Degradation System (MINDS), a multispecific nanobody-Fc chassis that co-engages multiple membrane receptors to promote highly efficient lysosomal co-depletion,, while also serving as a payload port. MINDS denotes a configurable scaffold with three interchangeable modules: (i) a highly efficient shuttle nanobody that binds a frequently internalized, non-competing epitope (preferably tumor-enriched) to drive endocytic routing; (ii) oncogenic-receptor nanobodies for multispecific co-engagement; (iii) defined conjugation sites for customizable payload attachment. Nanobodies (Nbs, ∼15 kDa) offer high affinity and stability, access to diverse including cryptic epitopes, and straightforward multivalent engineering with markedly improved potency and function^21–25^.

Using this chassis, we developed Tritazumab, which targets epidermal growth factor receptor (EGFR)^26^, hepatocyte growth factor receptor (HGFR/cMET)^27^, and transferrin receptor 1 (TfR1)^28^ to leverage their frequent co-overexpression in tumors for improved specificity and uptake. Tritazumab incorporates a non-transferrin-competing anti-TfR1 nanobody to initiate internalization through the endocytic activity of TfR1, and the construct was engineered to reliably induce lysosomal fusion to degrade all three receptors and efficiently release small-molecule payloads intracellularly. A Tritazumab-MMAE conjugate shows greater *in vitro* cytotoxicity than clinical comparators in matched models and *in vivo* tumor growth control in a lung cancer xenograft with favorable tolerability. Beyond chemotherapies, MINDS enables efficient delivery of intracellular degraders: a BRD4 molecular-glue conjugate improves the selectivity window in normal versus cancer cells, and an EZH2-targeting PROTAC (PROteolysis TArgeting Chimeras) conjugate achieves approximately 3-logs higher potency for intracellular degradation compared with the PROTAC alone^29^ *in vitro* in a model of lung cancer, and significantly reduces tumor volume *in vivo* in a model of breast cancer. We furthermore extended the platform to alternate tumor-associated antigen combinations including PD-L1, supporting the modularity and generalizability of MINDS. Together, these findings show that a single, modular scaffold can integrate surface receptor co-depletion with intracellular target control, addressing key limitations of monospecific ADCs.

## Results

### Discovery and Design of MINDS Targeting TfR1, EGFR and cMET

To characterize co-expression of oncogenic membrane receptors, we analyzed clinical proteomics (CPTAC) together with transcriptomics (MERAV) ^30,31^. Across tumor types, at least one or two of EGFR, cMET, and TfR1 were frequently elevated relative to matched normal tissues, with co-elevation of all three most evident in head and neck squamous cell carcinoma (HNSC) and clear cell renal cell carcinoma (CCRCC) (**Fig. S1**). Within non-small cell lung cancer (NSCLC), the squamous subtype (LSCC) showed upregulation of all three receptors, whereas lung adenocarcinoma (LUAD) exhibited increased EGFR and cMET but decreased TfR1. Complementary transcriptome analysis showed that EGFR, cMET, and TfR1 are each substantially higher in most NSCLC cell lines than in non-cancer lines (**Fig. S2**), supporting a multitarget strategy against these receptors.

We identified a repertoire of high-affinity Nbs to human TfR1from immunized llama serum using our proteomics-guided Nb discovery pipeline^32^. Among these, S11 emerged as a lead candidate, which binds strongly to the extracellular domain of TfR1 (residues 89-760) with K_D_ 313 pM by surface plasmon resonance (**Fig. 1A**). TfR1 consists of three subdomains: the apical, helical, and protease-like domains (**Fig. 1B**) ^33^. AlphaFold3 analysis of the S11-TfR1 complex yielded a high-confidence model with an interface predicted TM-score (ipTM) of 0.84 (**Fig. 1B**)^34^, consistent with near-experimental accuracy based on our recent benchmark study (Sang et al., *Science Advances*, accepted). The paratope is dominated by all three CDR loops, forming extensive contacts with TfR1 that likely underlie its high-affinity (**Fig. 1C**). S11 targets a distinct conformational epitope (aa 244-249, 351, 352, 354-357, 365, **Fig. 1B**) on the apical surface of the TfR1 homodimer, spatially separate from the transferrin (TF) binding site and absent from any reported TfR1-targeting biologics in the Protein Databank (**Fig. 1B, 1D**)^35^. Consistent with this structural prediction, competition assays confirmed that S11 does not interfere with TF binding, in contrast to another TfR1 Nb N4 (**Fig. 1E**). Because TF binds TfR1 with high affinity (low nM) and is present in serum at µM concentrations, biologics that compete with TF are likely to be outcompeted *in vivo*. This effect cannot be directly evaluated in standard mouse models due to the limited homology between mouse and human TfR1 (∼70%). Targeting a non-transferrin epitope therefore provides a favorable strategy for receptor-mediated uptake while better preserving physiological transferrin engagement and iron transport^28^.

**Figure 1.**
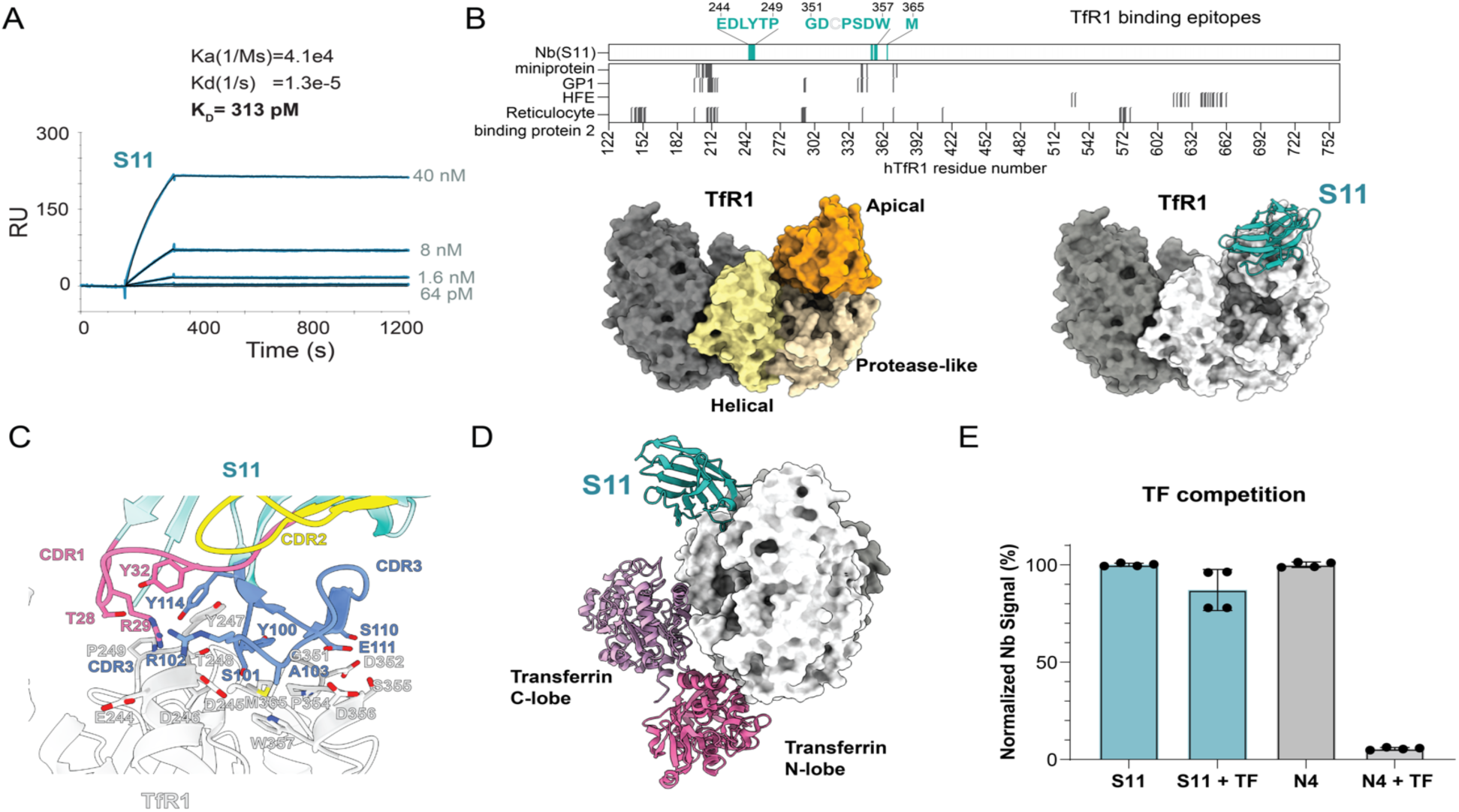
Discovery and characterization of TfR1 Nb S11. (A) Surface plasmon resonance (SPR) analysis of S11-TfR1 complex. S11 was immobilized on a CM5 sensor chip, and serial dilutions of ECD (extra cellular domain residues 89-760) of TfR1 were injected. Reference-subtracted traces (blue) are overlaid with a global 1:1 Langmuir fit (black), yielding K_D_ affinity of 313 pM. (B) High-confidence structural model (ipTM= 0.84) of the TfR1-S11 complex predicted by Alphafold-3. Top panel: epitope residues of S11 (blue) and other TfR1 binders (light grey) including miniprotein (PDB: 6WRX), Machupo GP1(PDB: 3KAS), HFE (PDB: 1DE4), and Reticulocyte binding protein 2 (PDB: 6D04). Bottom left: three TfR1 domains (apical, helical, and protease-like) on the dimeric TfR1 were shown. Bottom right: the predicted TfR1-S11 complex. (C) Modeled interaction. CDRs on S11 are colored: CDR1 (pink), CDR2 (yellow) and CDR3 (blue). Interface residues are labeled. (D) Superposition of AF3-modeled S11 and transferrin (TF) (PDB:1SUV)^30^ on the TfR1. (E) Competitive ELISA assay of S11 and a control Nb N4 that we isolated, which competes with TF, in the presence of excess TF (N = 4).

In parallel, we identified Nb15, a strong binder to the cMET ectodomain (**Fig. S3**) and incorporated the established EGFR Nb 7D12^36^. We then fused S11 (TfR1), 7D12 (EGFR), and Nb15 (cMET) to the N-terminus of a human IgG1 Fc to create Tritazumab, a trispecific Nb-Fc degrader (**Fig. 2A-B**). S11 was positioned at the distal N-terminus to help accessibility to the dimeric TfR1, whereas 7D12 and Nb15 were incorporated using flexible linkers (GGGGS)_5_ to preserve binding to EGFR and cMET (**Fig. 2C**). The IgG1-Fc scaffold was included to enhance avidity and as a design rationale to improve *in vivo* half-life^37,38^, although the specific pharmacokinetic properties of Tritazumab were not directly evaluated in this study. Simultaneous targeting of TfR1 together with oncogenic receptors would promote receptor-mediated internalization and lysosomal trafficking while broadening receptor coverage in tumor cells. This architecture also provides a modular platform for conjugation of diverse therapeutic payloads.

**Figure 2.**
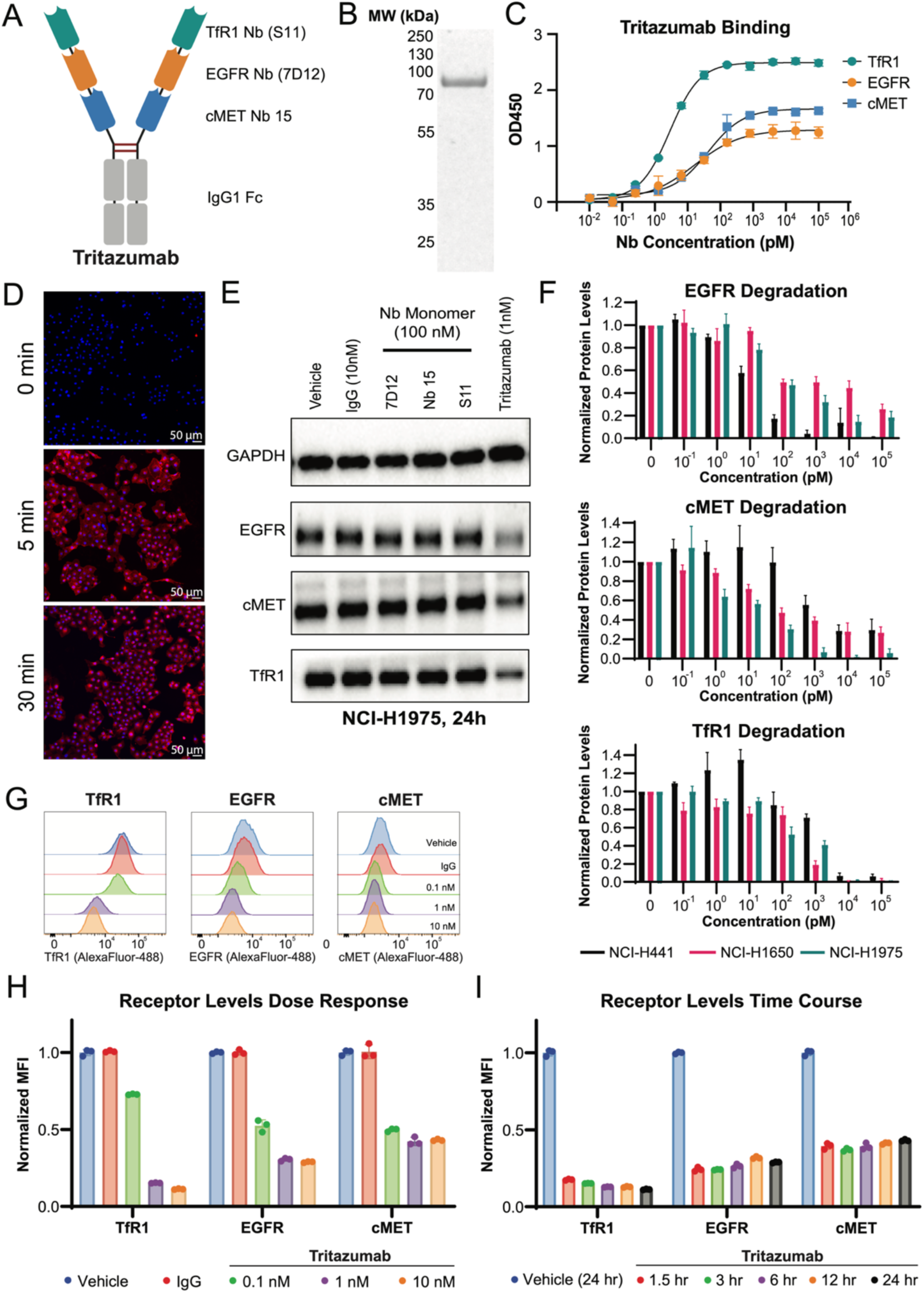
Discovery and engineering of Tritazumab. (A) Design architecture of Tritazumab. (B) SDS-page gel of Tritazumab. (C) Target engagement assessment of Tritazumab at various concentration by ELISA assays (N = 3). (D) Visualization of endocytosis by Tritazumab (10 nM) conjugated with Alexa Fluor-647 dye, in NCI-H1975 cells at 0, 5, and 30 mins by confocal microscopy. Results shown are representative of two independent experiments. (E) Western blotting (WB) analysis of EGFR, cMET and TfR1 in NCI-H1975 cells treated with Tritazumab (1 nM), vehicle, IgG (10 nM), 7D12 (100 nM), Nb15 (100 nM), or S11(100 nM) for 24h (N = 2). Total cell lysates were used for WB and GAPDH was used as a loading control. WB results shown are representative of 2 independent experiments. (F) Quantification of WB concentration-dependent binding of EGFR, cMET and TfR1 by Tritazumab in NCI-H441, NCI-H1650 and NCI-H1975 cells at different concentrations after 24h treatment (from Fig. S4B-D). Results shown are representative of three independent experiments. Total cell lysates were used for WB and GAPDH was used to normalize the protein levels. (G) Representative flow cytometry assessment of cell-surface EGFR, cMET and TfR1 degradation by Tritazumab at 0.1, 1 and 10 nM concentration after 24h compared to vehicle and IgG (10 nM) in NCI-H1975 cells (N = 3). NCI-H1975 cells were treated with Tritazumab at the indicated concentrations with vehicle and IgG as controls for 24h, followed by detection using AF488 -conjugated monomer-Nb-Fc constructs. (H) Quantification of cell-surface EGFR, cMET and TfR1 protein levels from panel G and its replicates. (I) Quantification of cell-surface EGFR, cMET and TfR1 protein levels in NCI-H1975 cells treated with Tritazumab (10 nM) at indicated times (N = 3).

### Cellular Uptake and Target Degradation in NSCLC Cells

We first evaluated the cellular uptake of Tritazumab using a NSCLC model cell line NCI-H1975. Intracellular trafficking of fluorophore-conjugated (Alexa Fluor-647) Tritazumab (10 nM) was monitored by confocal fluorescence microscopy. Strong fluorescence signal was observed by 5 minutes of incubation, consistent with rapid cell association and uptake (**Fig. 2D**). To assess functional degradation, we treated NCI-H1975 cells with 1 nM Tritazumab for 24 h and compared target levels to cells treated with isotype IgG or monomeric Nbs. Immunoblotting revealed substantial degradation (>50%) of EGFR, cMET, and TfR1, whereas control treatments had no measurable effects (**Fig. 2E**).

Next, we examined Tritazumab across multiple NSCLC cell lines with variable expression profiles of EGFR, cMET and TfR1. NCI-H441, NCI-H1650 and NCI-H1975 all have elevated *EGFR* expression and marked upregulation of *MET* and *TFRC* compared to normal lung fibroblasts (**Fig. S4A**). These cell lines also differ in EGFR mutational status and MET activity, providing a genetically diverse panel for evaluating degradation activity ^39^.

Across NSCLC cell lines, Tritazumab induced potent receptor degradation (**Fig. 2F, S4B-G**). In NCI-H1975, which harbors EGFR-L858R/T790M mutations and has high cMET activity^39^, Tritazumab degraded EGFR, cMET, and TfR1 with DC_50_ values of 52.4 ± 8.2, 38.2 ± 11.1, and 351 ± 9.2 pM, respectively (**Fig. 2F, S4B**). In NCI-H441 cells, which overexpress EGFR-WT and MET, the corresponding DC_50_s were 10.3 ± 4.1, 494 ± 22.7 and 1044 ± 236 pM, respectively (**Fig. 2F, S4C**). In NCI-H1650 cells, which harbor an EGFR-exon19del and low MET activity^39^, the DC_50_s were 68.8 ± 8.5 pM for EGFR, 30.9 ± 8.2 pM for cMET and 482 ± 28.3 pM for TfR1 (**Fig. 2F, S4D**). Flow cytometry analysis further showed concentration-dependent manners in surface EGFR, MET, and TfR1 in H1975 and H1650 cells (**Fig. 2G-H, S4E-G**). Near-maximal surface loss was observed within 1.5 h and remained sustained after 24 h of treatment (**Fig. 2I**).Collectively, these data show that Tritazumab drives fast, potent and sustained multi-receptor degradation across genetically diverse NSCLC models.

### Mechanism of potent membrane protein degradation

We compared Tritazumab with monospecific constructs including 7D12-Fc, Nb15-Fc, and S11-Fc, in NCI-H1975 cells. Whereas the EGFR-targeting construct 7D12-Fc did not reduce EGFR levels, the cMET-targeting Nb15-Fc and TfR1-binding S11-Fc each produced moderate, target-restricted degradation (**Fig. 3A**). In contrast, Tritazumab triggered robust co-degradation of EGFR, MET, and TfR1 at 1 nM (the lowest tested concentration), consistent with enhanced degradation from trispecific engagement (**Fig. 3A**). Flow cytometry binding showed that Tritazumab and the monospecific controls displayed similarly low EC_50_s of approximately 100 pM in NCI-H1975 cells, whereas Tritazumab reached a considerably higher maximal binding signal (**Fig. 3B**). This observation implies greater cumulative receptor engagement on the same cells, which may contribute to more efficient intracellular uptake. Tritazumab also displayed a higher maximal binding signal in NCI-H1975 cells than healthy HFL-1 control cells (**Fig. S7**), indicating a potential selectivity advantage based on total receptor engagement.

**Figure 3.**
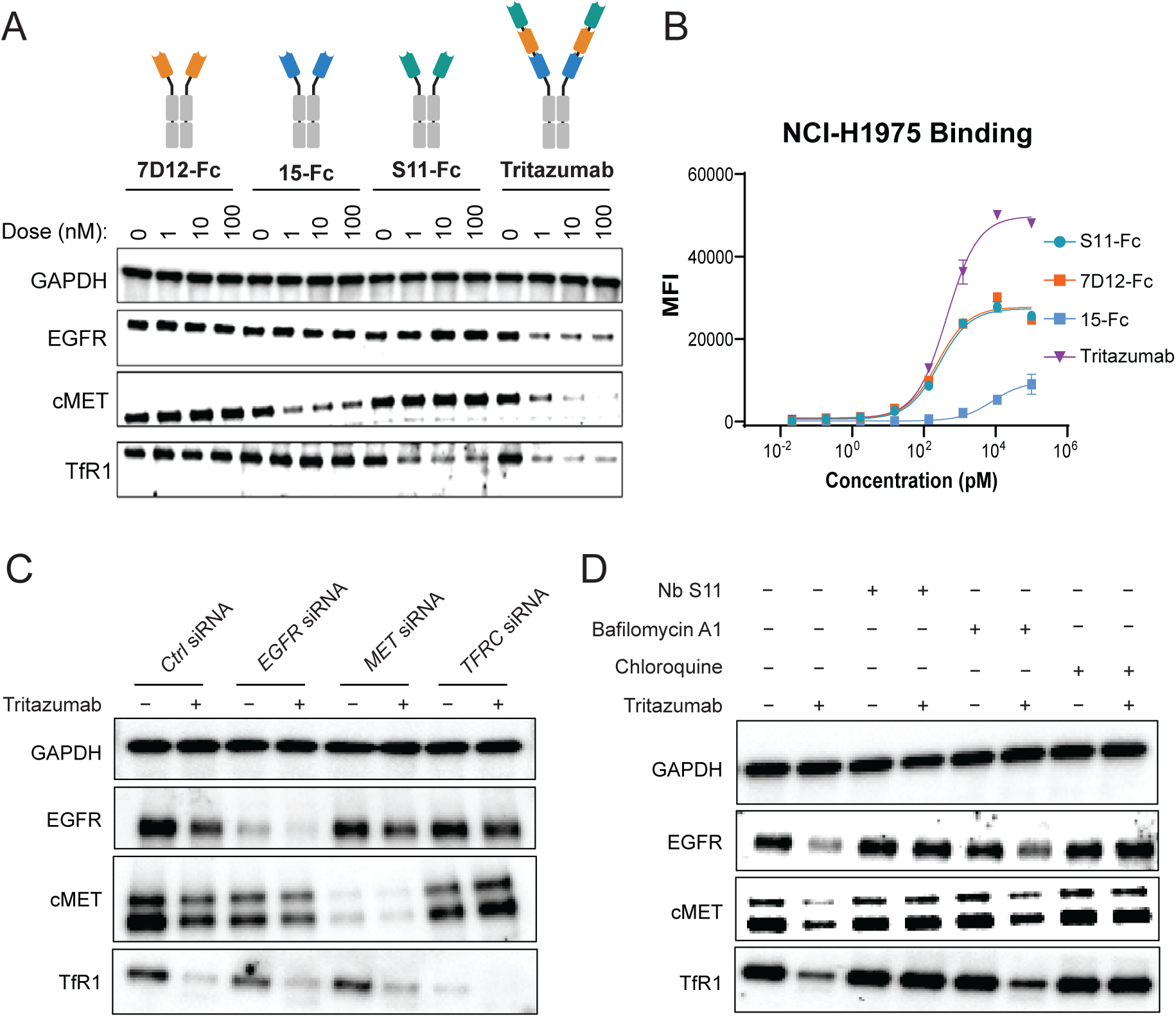
Mechanism of membrane protein degradation induced by Tritazumab. (A) WB results of EGFR, cMET and TfR1 degradation by 7D12, Nb15 and S11-monomer-Nb-Fc construct compared to Tritazumab in NCI-H1975 cells at the indicated concentration after 24h treatment (N = 2). Total cell lysate was used for WB and GAPDH was used a loading control. (B) Flow cytometry cell-binding affinity of 7D12, Nb15 and S11-monomer-Nb-Fc constructs compared to Tritazumab in NCI-H1975 cell line (N = 2). (C) Representative WB results of EGFR, cMET and TfR degradation by Tritazumab at 10 nM for 24h treatment upon *EGFR*, *MET* and *TFRC* knockdown by siRNA in NCI-H1975 cells (N = 2). Total cell lysate was used for WB and GAPDH was used a loading control. (D) Representative WB results of EGFR, cMET and TfR1 degradation rescue after pre-treatment of NCI-H1975 cells with 100 nM of S11, 50 nM of Bafilomycin A1 or 100 μM of chloroquine for 4h, followed by 10 nM of Tritazumab treatment for 18h (N = 2). Total cell lysate was used for WB and GAPDH was used a loading control.

To determine which receptor predominantly drives co-degradation, we used siRNA to individually knock down *EGFR*, *MET*, or *TFRC* in NCI-H1975 cells. Notably, only TfR1 knockdown rescued receptor levels following Tritazumab treatment (**Fig. 3C**), indicating that TfR1 engagement is the primary driver of Tritazumab-induced endocytosis^40^. Next, we pretreated cells with either Nb S11, chloroquine (a lysosomal acidification inhibitor), or bafilomycin A1 (an autophagy inhibitor). Nb S11 and chloroquine completely abrogated Tritazumab-induced degradation of EGFR, cMET, and TfR1, while bafilomycin A1 showed partial rescue (**Fig. 3D**).

To test whether TfR1-driven co-degradation could be extended to an alternative target combination, we generated a second trispecific construct, S11-7D12-KN035-Fc, targeting TfR1, EGFR, and PD-L1. This construct likewise induced robust degradation of EGFR, PD-L1, and TfR1 in NCI-H1975 cells (**Fig. S5**), supporting the modularity of the MINDS platform and the broader generalizability of TfR1-directed multispecific degradation.

Together, the TFRC knockdown-rescue experiment, the matched bispecific controls, and an additional TfR1-based trispecific construct targeting EGFR and PD-L1 support TfR1 as the dominant uptake in the tested MINDS constructs, while the precise relative contributions of receptor abundance, arm affinity, and intrinsic trafficking remain to be defined.

### Development of Tritazumab-MMAE

We generated a Tritazumab-MMAE conjugate to evaluate targeted payload delivery using monomethyl auristatin E (MMAE)^8^. Conjugation used cysteine-maleimide chemistry with a lysosomally cleavable valine-citrulline (VC)-MMAE linker-payload, which is the payload component of clinical vedotin ADCs. Accessible thiols in the IgG1-Fc hinge of the Tritazumab scaffold were conjugated following controlled partial reduction. Cysteine C103, normally involved in light-chain pairing but surface-exposed with mild reduction in the MINDS architecture, served as a primary conjugation site (**Fig. 4A**). Additional conjugation could be enabled at the “CPPC” motif following stronger reduction conditions with TCEP if needed. Using controlled cycsteine reduction to mitigate potential perturbation of the scaffold, we generated Tritazumab ADCs with a drug-antibody ratio (DAR) of ∼ 2, as determined by HPLC coupled to quadrupole time-of-flight mass-spectrometry (Q-TOF MS) analysis (**Fig. 4B**). ELISA further confirmed that Tritazumab-MMAE conjugate retained similar binding to EGFR, cMET and TfR1 comparable to that of unconjugated Tritazumab (**Fig. 4C-E**).

**Figure 4.**
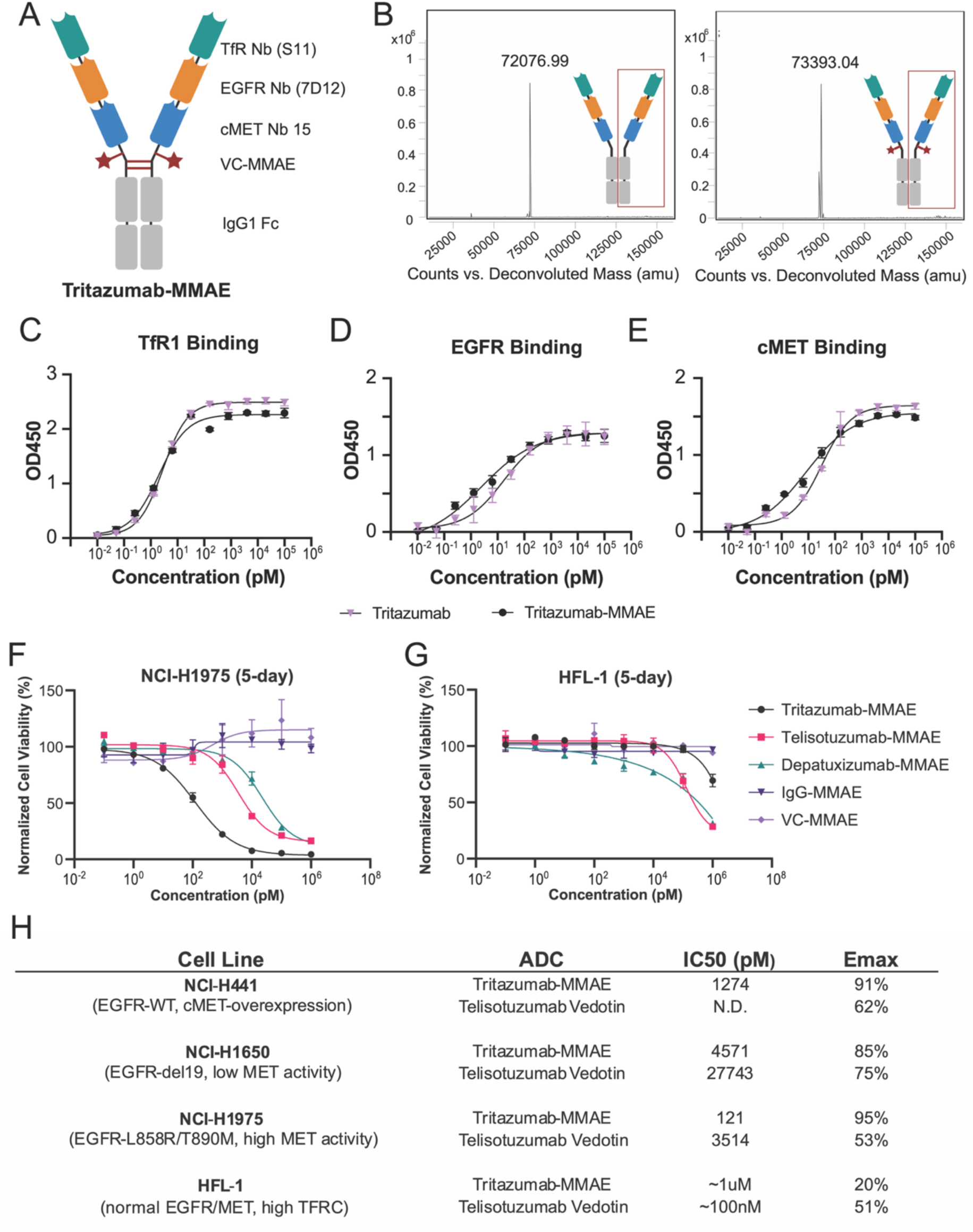
Development of the Tritazumab-MMAE ADC. (A) Architecture of Tritazumab highlighting accessible cysteine residues for MMAE conjugation. (B) Q-TOF assessment of Tritazumab compared to Tritazumab-MMAE. (C-D) Receptor binding assessment of Tritazumab vs. Tritazumab-MMAE against (C) TfR1, (D) EGFR and (E) cMET via ELISA binding assay. (F-G) Cell viability of Tritazumab-MMAE, Telisotuzumab Vedotin, Depatuxizumab-MMAE, IgG-MMAE and VC-MMAE in (F) NCI-H1975 and (G) HFL-1 cells. The cells were treated with vehicle or the indicated compounds at the indicated concentrations for 5 days. The mean IC_50_ value ± SD for each concentration point (in technical duplicates from N = 3 biological experiments) is shown in the curves. (H) Table summary of Tritazumab-MMAE vs Telisotuzumab Vedotin in other NSCLC cell lines.

The cytotoxic efficacy of Tritazumab-MMAE was compared to two ADC comparators in NSCLC cell lines: the FDA approved cMET-targeting Telisotuzumab-MMAE (Telisotuzumab Vedotin) and an EGFR-targeting Depatuxizumab-MMAE^41,42^. In NCI-H1975 cells, Tritazumab-MMAE (IC_50_ = 120.8 ± 9.4 pM) displayed 30-fold more potent cell killing than Telisotuzumab-MMAE (IC_50_ = 3,514 ± 85.9 pM) and 183-fold more potent cell killing than Depatuxizumab-MMAE (IC_50_ = 22,095 ± 86.7 pM). (**Fig. 4F**).

Notably, Tritazumab-MMAE maintained sub-nM potencies and >85% maximal inhibition (I_max_) across NSCLC lines with varied cMET activities, including high cMET activity NCI-H441 cells (IC_50_ = 1274 ± 166.5 pM) and low cMET activity NCI-H1650 cells (IC_50_ = 4571± 345.7 pM) (**Fig. 4H, S6**). Moreover, Tritazumab-MMAE showed limited activity against a normal human lung fibroblast (HFL-1) and human keratinocyte (HaCat) cell lines, with IC_50_ > 1μM, indicating a favorable *in-vitro* selectivity window (**Fig. 4G, S8D**). This favorable therapeutic window likely reflects the low cumulative expression of EGFR, cMET, and TfR1 in normal tissues compared to cancer cells (**Fig. S4**) resulting in lower volume of antibody engagement (**Fig. S7**), which restricts Tritazumab binding volume and payload internalization in healthy cells.

### Contribution of trispecific architecture to Tritazumab activity

To define how trispecificity contributes to the activity of Tritazumab, we compared it against bispecific controls in degradation, binding and cell-killing studies. We utilized the same design rationale with anti-TfR1 Nb S11 at the N-terminus followed by a (GGGGS)_5_ linker and anti-EGFR Nb 7D12 (S11-7D12-Fc) or anti-cMET Nb15 (S11-Nb15-Fc) (**Fig. S8A**). Bispecific constructs show comparably high expression and purity to Tritazumab (data not shown), consistent with the suitability of affinity-matured Nbs as building blocks for multivalent engineering^24,25,43^.

Both constructs induced degradation of their respective target combinations, whereas Tritazumab maintained efficient degradation across all three receptors (**Fig. S8B**). Thus, although the bispecific formats retained strong activity within narrower target combinations, the trispecific architecture expanded degradation breadth to a broader receptor set relevant to tumor heterogeneity. Tritazumab-MMAE showed significantly greater antiproliferative activity than either bispecific-MMAE conjugate in NCI-H1975 cells at 1 nM (**Fig. S8C**).

### *In-vivo* efficacy and safety of Tritazumab-MMAE

We assessed *in vivo* efficacy and safety of Tritazumab-MMAE in an NCI-H1650 human NSCLC xenograft. We prioritized the NCI-H1650 cell line, which harbors the EGFR-exon 19 deletion, a frequently observed mutation in NSCLC patients with low MET activity^39,44,45^. Notably, clinical Telisotuzumab (cMET)-MMAE showed minimal cell killing efficacy in NCI-H1650^46^. The strong *in-vitro* potency of Tritazumab-MMAE prompted us to test whether it could suppress tumor growth *in vivo*.

Mice bearing *subcutaneous* flank tumors (injected with 5 x 10^6^ NCI-H1650 cells) were randomized to vehicle or Tritazumab-MMAE and dosed 1 mg kg⁻¹ intraperitoneally (i.p.) on days 1 and 12 (**Fig. 5A**). Tumor volumes and body weights were monitored every other day up to day 21 (n=6/arm), with blood collected from a subset for hematology (n=3/arm). Tritazumab-MMAE produced marked and significant tumor growth suppression beginning after the second dose, achieving ∼ 90 % tumor growth inhibition at day 21 relative to vehicle control (**Fig. 5B-C**). Following Tritazumab-MMAE treatment, we observed the tumor-bearing mice gradually gained weight with no evidence of overt toxicity, and by study end (day 21) body weight was higher in Tritazumab-MMAE treated mice (**Fig. 5D**). Hematology parameters showed no significant differences (**Fig. 5E**). Together with the *in vitro* cytotoxicity results, these data support the antitumor activity and general tolerability of Tritazumab-MMAE in the low-*MET* model.

**Figure 5.**
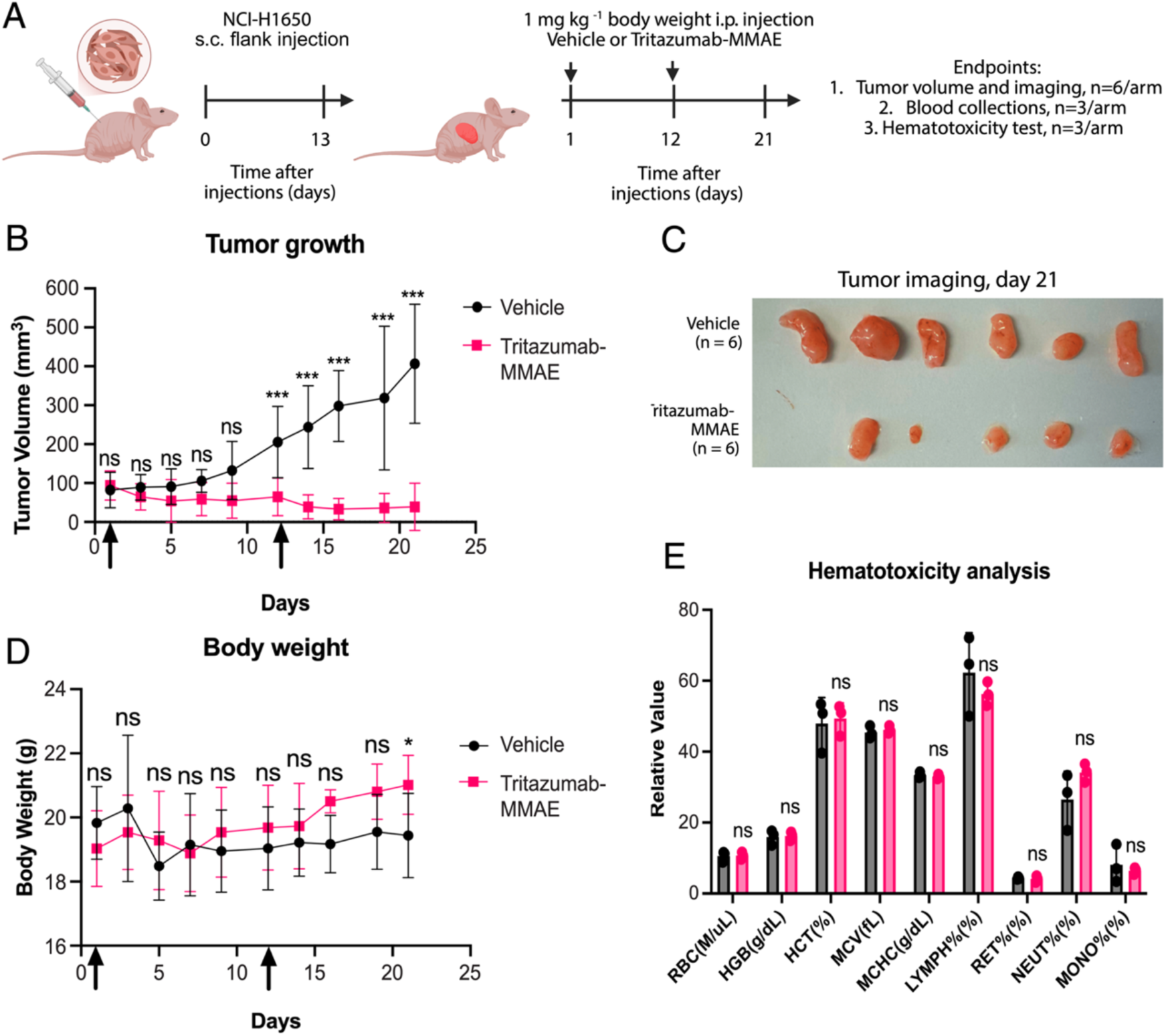
*In vivo* evaluation of the Tritazumab-MMAE ADC. (A) Schematic illustrating the mouse experiments to assess the safety and efficacy of Tritazumab-MMAE. (B) Average tumor growth curves showing Tritazumab-MMAE led to significant tumor inhibition compared to vehicle groups. Data are presented as median values with standard errors of the mean from n = 6 mice for both treatment and control groups. P-values (ns - not significant, or *** - p-value<0.001) were determined by unpaired two-tailed t-tests. (C) Images of harvested tumors on day 21. (D) Average weight measurement showing Tritazumab-MMAE led to increased body weight compared to vehicle groups on day 21. P-values (ns - not significant, or * - p-value<0.05) were determined by unpaired two-tailed t-tests. (E) Hematotoxicity assessment following second injection of Tritazumab-MMAE showing no toxicities. The analysis was performed using mice blood samples harvested from both treatment or vehicle group (n=3) at day 21 and were determined by unpaired two-tailed t-tests.

### Combined membrane and intracellular protein degradation

To extend MINDS beyond traditional cytotoxic payloads, we conjugated intracellular degraders to Tritazumab (**Fig. 6A**), including a BRD4 molecular glue (MG) (**Fig. 6B**) and an EZH2-targeting PROTAC (**Fig. S11A**), using the same VC linker and cysteine-maleimide chemistry.

**Figure 6.**
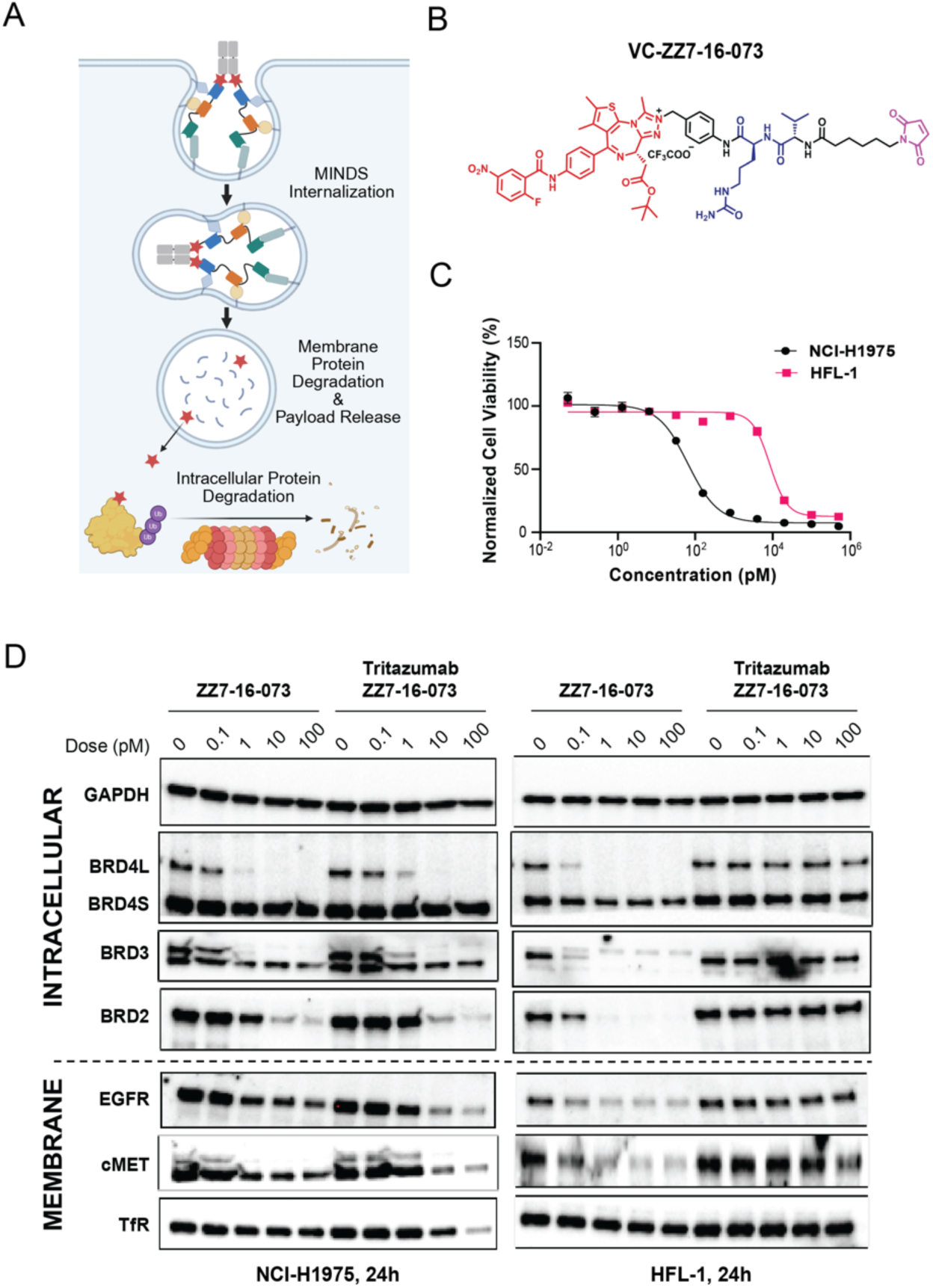
Development of membrane and intracellular protein degradation modality. (A) Schematic of our bi-directional system to degrade membrane and intracellular proteins. (B) Chemical structure of VC-ZZ7-16- 073. (C) Cell viability of Tritazumab-ZZ7-16-073 in NCI-H1975 and HFL-1 cells. The cells were treated with Tritazumab-ZZ7-16-073 at the indicated concentrations for 5 days. The mean value ± SD for each concentration point (in technical duplicates from N=3 biological experiments) is shown in the curves. (D) WB result of ZZ7-16-073 and Tritazumab-ZZ7-16-073 at 0, 0.1, 1, 10 and 100 pM treatments in degrading membrane and intracellular proteins in NCI-1975 and HFL-1 cells (N = 2). Total cell lysate was used for WB and GAPDH was used a loading control.

BRD4 is among the most actively pursued therapeutic targets, with numerous BET inhibitors evaluated clinically, but single-agent dosing has been limited by on-target hematologic toxicity ^47^. The BRD4 MG ZZ7-16-073 recruits DCAF16 and potently degrades BRD4 and other BET proteins in cells^48^. We synthesized VC-ZZ7-16-073 (**Methods**) and generated Tritazumab-ZZ7-16-073 with a DAR of ∼ 2, confirmed by Q-TOF MS (**Fig. S9**).

MG ZZ7-16-073 displayed minimal therapeutic window, inducing comparable cytotoxicity in NCI-H1975 cancer cells and HFL-1 normal fibroblasts (**Fig. S10A**). In contrast, Tritazumab-ZZ7-16-073 maintained strong BRD4-L degradation activity in NCI-H1975 cells (**Fig. 6D**) and MDA-MB-231 cells (**Fig. S10B**), while showing no detectable BRD4-L degradation in HFL-1 cells (**Fig. 6D**). This pattern aligns with the lower cumulative expression of EGFR, MET, and TfR1 in HFL-1 cells (**Fig. S4A**), which may limit Tritazumab binding and cellular uptake. Correspondingly, Tritazumab-ZZ7-16-073 displayed a marked *in vitro* selectivity window, with cytotoxicity in NCI-H1975 cells (IC₅₀ = 65 ± 9.8 pM) compared with HFL-1 cells (IC_50_ = 8,425 ± 38.2 pM), representing a two-log therapeutic window. In contrast, free ZZ7-16-073 induced highly potent cytotoxicity in HFL-1 cells (femtomolar IC_50_; **Fig. S10A**), implying a substantially improved safety margin upon Tritazumab conjugation.

We evaluated the *in vivo* efficacy of Tritazumab-ZZ7-16-073 in a triple negative breast cancer (TNBC) xenograft model using MDA-MB-231 cells (**Fig. S10C-E**). Although EGFR and cMET expression levels in this model are lower than in other TNBC lines (such as MDA-MB-468)^49^, Tritazumab-ZZ7-16-073 treatment produced pronounced tumor growth suppression. Mice bearing mammary fat pad tumors were randomized to vehicle or Tritazumab-ZZ7-16-073 and treated intraperitoneally at 1 mg kg⁻¹ on days 20, 23, and 27 (n = 8 per group). Tumor growth was monitored until day 30. Tritazumab-ZZ7-16-073 achieved ∼74 % tumor growth inhibition relative to vehicle controls without evidence of overt toxicity (**Fig. S10C,D**).

We next targeted EZH2, a lysine methyltransferase strongly implicated in multiple cancers, using the cereblon-recruiting PROTAC MS177^29^. We prepared VC-MS177 and conjugated it to Tritazumab to yield Tritazumab-MS177 (DAR2 based on Q-TOF analysis) (**Fig. S11 B, C**). Tritazumab-MS177 degraded EZH2 at 1 nM in the MDA-MB-468 (breast cancer) cells, about 1,000-fold more potent than the free MS177 (1 μM) and led to a stronger reduction of the downstream chromatin mark H3K27me3 (**Fig. S11 D**).

Together, our results demonstrate that the MINDS chassis can coordinate membrane receptor co-degradation with intracellular protein degradation. Conjugation of Tritazumab to a BRD4 molecular glue improved the selectivity window relative to the free compound, whereas conjugation to an EZH2 PROTAC markedly enhanced intracellular degradation potency. These findings indicate that receptor-directed delivery via MINDS can extend targeted degradation strategies to intracellular proteins while maintaining tumor-selective activity.

## Discussion

Despite transformative clinical impact, ADCs that rely on a single receptor remain limited by heterogeneous antigen expression, compensatory signaling, and variable internalization kinetics. The MINDS chassis addresses these constraints with a compact Nb-Fc architecture that co-engages multiple tumor-associated receptors and converts that engagement into rapid lysosomal trafficking and efficient payload delivery. Tritazumab, which targets TfR1, EGFR, and MET, achieves rapid, sub-nM receptor degradation in NSCLC models and, when conjugated to MMAE, translates uptake into potent antitumor activity while sparing normal lung fibroblasts. Nbs furnish a compact, robust scaffold, high-affinity binding, access to diverse and cryptic epitopes, and modularity that supports advanced multivalent engineering and production. Broadly, MINDS functions as a convergent modality that unifies multi-receptor degradation, delivery of cytotoxic payloads and intracellular degraders within one platform.

A central design principle is the use of TfR1 as a noncompeting internalization shuttle ^28^. Our high-affinity S11 Nb recognizes a novel epitope that does not interfere with transferrin binding, permitting frequent endocytosis without perturbing iron metabolism. Consistent with this mechanism, TFRC knockdown rescues receptor levels after Tritazumab treatment and lysosomal inhibition attenuates degradation, indicating predominant routing to lysosomes through TfR1 engagement^40,50^. More generally, the shuttle epitope should be preferentially displayed in tumors, frequently internalized, and nonoverlapping with essential ligand sites to reduce uptake in safety-critical tissues. Judicious selection of such a shuttle together with oncogenic receptors provides a mechanistic basis for coordinated multi-receptor co-degradation.

Multispecificity and high valency appear central to both degradation efficiency and functional selectivity. Although cell bindings are comparable between monospecific controls and trispecific design, Tritazumab exhibits higher maximal binding, consistent with greater net occupancy and potential receptor clustering. Productive membrane receptor engagement, and ligand-induced internalization and trafficking, may require simultaneous engagement of two or more receptors in close proximity and at sufficient surface density^51,52^; however, the biology of these processes remains to be understood and warrants further investigation. Tumor cells more likely to meet this threshold because they frequently co-express multiple receptors at elevated levels ^53–55^, which raises the bar for stable binding, uptake, and payload release on normal tissues and thereby improves on-target and off-tumor specificity.

These mechanistic features translate into head-to-head readouts. Tritazumab-MMAE maintains binding to all three targets, shows higher *in-vitro* potency than two clinical ADCs, and despite a low dose (1 mg/kg), suppresses tumor growth in a NSCLC xenograft with tolerability under the study conditions. The same chassis also enables intracellular target control: a BRD4 molecular-glue conjugate improves *in-vitro* therapeutic window by > 100-fold compared to the free compound, while preserving pM activity in cancer cells, and an EZH2-targeting PROTAC conjugate achieves a marked 1,000-fold improvement for intracellular degradation in a breast cancer model. *In vivo,* Tritazumab conjugated to a BRD4 molecular glue showed significant reduction in tumor volume without observable toxicity in a xenograft model of breast cancer, further validating the efficacy observed *in vitro*. These catalytic applications are dose sparing and expand target space^56–58^.

Our study has several limitations. First, the breadth of the platform is currently supported by two trispecific proof-of-concept designs, and more extensive characterization of DAR distributions, plasma stability, and related developability features will be important for future development. Even so, the present data support a generalizable multispecific strategy in which a cancer-preferential, frequently internalized shuttle epitope is combined with disease-relevant receptors to establish a composite antigen threshold on tumor cells and coupled to a payload-linker system suited to the underlying trafficking route. Second, although Nbs have shown favorable clinical safety profiles, humanization using dedicated tools such as *Llamanade* may help mitigate potential immunogenicity^59,60^. Third, although we observed marked tumor selectivity and tolerability *in vitro* and in xenograft mouse models, TfR1 is broadly expressed in normal tissues. Third, although we observed marked tumor selectivity and tolerability *in vitro* and in xenograft mouse models, TfR1 is broadly expressed in normal tissues. In addition, specific pharmacokinetic parameters of Tritazumab, including clearance, half-life, and volume of distribution, were not determined in the present study and will require direct evaluation. Accordingly, formal dose-dependent biodistribution, pharmacokinetic, and toxicology studies will be required, ideally in pharmacologically relevant species such as non-human primates, whose TfR1 more closely resembles the human receptor than mouse TfR1.

## Methods

### Cell Lines, Tissue Culture and Transfection

NCI-H441 (CRM-HTB-174), NCI-H1650 (CRL-5883), NCI-1975 (CRL-5908), MDA-MB-468 (HTB-132) and HFL1 (CCL-153) were purchased from the American Tissue Culture Collection (ATCC, Manassas, VA). NCI-H441, NCI-H1650, NCI-1975 and MDA-MB-468 cells were cultured in RPMI 1640 medium (Thermo Fisher Scientific Inc., Waltham, Massachusetts) supplemented with 10% heat-inactivated fetal bovine serum GibcoTM (FBS) (Life Technologies, Grand Island, NY) and 1% GibcoTM Penicillin/Streptomycin (Life Technologies, Grand Island, NY). HFL1 cells were cultured in Ham’s F-12K (Kaighn’s) Medium (Thermo Fisher Scientific Inc., Waltham, Massachusetts) supplemented with 10% heat-inactivated FBS and 1% Penicillin/Streptomycin. All cells were incubated at 37 °C in a standard humidified incubator containing 5% CO2 and 95% O2.

### Antibodies and Immunoblotting

Total cell lysate was used for western blots, as previously described.^52^ The following primary antibodies were used in the study: Vinculin (Cell Signaling Technology [CST], 13901), GAPDH (14C10) (CST, 2118), EGF Receptor (D38B1) (CST, 4267), cMET (3D4) (Thermo Fisher Scientific Inc., Cat#37-0100) and TfR (Thermo Fisher Scientific Inc., Cat#13-6800). Blots were imaged using fluorescence-labeled secondary antibodies on LI-COR Odyssey CLx Imaging Systems (LI-COR Biosciences, Lincoln, NE) or iBright^TM^ CL750 Imaging Systems (Thermo Fisher Scientific Inc., Waltham, MA).

### Cell Viability Assay

The cell viability assay was performed as previously described.^52^ Briefly 5 x 10^4^ cells were seeded per well into 96-well microplates. After 24h, cells were treated with 10-fold serially diluted compounds in duplicates for indicated days. Cell viability was evaluated using WST-8 reagent (CK04, Dojindo). Absorbance signals were obtained with Infinite F PLEX plate reader (TECAN, Morrisville, NC) at 450 nm with 690 nm as reference wavelength after 3 h incubation at 37 °C. GraphPad Prism 8 was used in the analysis of GI_50_ values from the data of 2-3 independent experiments.

### Purification of Tritazumab

Plasmids for Nb-IgG Fc fusion constructs were synthesized from Synbio technologies using a propriety IgG Fc vector. All antibody constructs were transiently transfected into Expi293F cells using Expifectamine 293 kit system (Gibco, Thermo Fisher Scientific Inc., Waltham, MA) according to manufacturer protocols. Expifectamine enhancers (Gibco, Thermo Fisher Scientific Inc., Waltham, MA) were added after 20 hours to improve protein expression. 5 days after transfection, expression cell culture media was collected and centrifuged at 300xg for 5 minutes to isolate the supernatant containing secreted Nb-Fc. Supernatants were incubated with recombinant Protein A resin (Marvelgent) for 1 hour at 4°C and washed with cell culture grade PBS. Immobilized Nb-Fc constructs were eluted with pH 3.0 glycine buffer and immediately brought to a neutral pH with pH 8 1M Tris solution. Nb-Fc fusion constructs were further purified with size exclusion chromatography using a Superdex 75 Increase 10/300GL column installed in an Akta Pure FPLC system. Purified Nb-Fc constructs were purified of endotoxins using Detoxi-Gel endotoxin removing gel (Thermo Fisher Scientific Inc., Waltham, MA) and confirmed of endotoxin levels using an LAL assay (Genscript ToxinSensor, Nanjing, China).

### Generation of Tritazumab-ADC

Purified endotoxin free Nb-Fc constructs were diluted to 1.05mg/ml and incubated with 20x molar excess of TCEP for 1 hour at 25°C. Reduced Nb-Fc constructs were buffer exchanged with several rounds of concentration and dilution using a 30kDa cut-off concentrator. Nb-Fc constructs were diluted to 1.05mg/ml and mixed with 20x molar excess of small molecule payloads with reactive maleimide moieties (5% v/v DMSO final). Cysteine-maleimide reactions ensued for 16 hours at 25°C. Excess unconjugated small molecules were removed using several rounds of concentration and dilution using a 30kDa cut-off concentrator. Conjugation efficiency and DAR measurements were confirmed using SDS-PAGE and Q-TOF mass spectrometry.

### Protein Q-TOF Analysis

Antibody samples were dialyzed into ammonium acetate buffer and analyzed by an Agilent 1200 series system with DAD detector and 2.1 mm x 150 mm Zorbax 300SB-C18 5 μ column for chromatography. Samples (20 μL, ∼ 1 - 5µM) were injected onto a C18 column at room temperature with a flow rate of 0.4 mL/min. Chromatography was performed with the solvent as follows: water containing 0.1% formic acid and 3% acetonitrile was designated as Solvent A while acetonitrile containing 0.1% formic acid was designated as Solvent B. For the reduced antibody mass measurements, the proteins were first treated with TCEP, and the linear gradient was set such that 5% B from 0 - 1 min, 5 - 70% B from 1 - 6 min, 70% B from 6 - 8 min, and 5% B from 8 - 12 min. For the intact antibody mass measurements, the linear gradient was set such that 5% B from 0 - 1 min, 5 - 70% B from 1 - 6 min, 70% B from 6 - 8 min, 70 - 80 % B from 8 - 8.5 min and 80% B from 8.5 - 12 min. Mass spectra data was acquired in positive ion mode using an Agilent 6320 TOF with an electrospray ionization (ESI) source. The data was analyzed by Agilent MassHunter BioConfirm 12.0.

### ELISA Binding

Relative binding affinities of Nb constructs were assessed using indirect ELISA experiments. A 96-well ELISA plate (R&D system, Minneapolis, MN) was coated with recombinant antigens of interest (cMET, TfR1, and EGFR) overnight at 4°C in a coating buffer (15 mM sodium carbonate, 35 mM sodium bicarbonate, pH 9.6). ELISA plates were washed with PBST (PBS + 0.05% Tween20) and blocked (PBST + 5% milk powder) for 2 hours at room temperature. Nb or Nb-Fc constructs were serially diluted 5-fold in blocking buffer and added to the blocked plates and incubated for 2 hours at room temperature.

For non-Fc fused Nb constructs, HRP conjugated anti-T7 polyclonal antibodies (Invitrogen) were added at 1:7,500 dilutions. For Fc-fused Nb constructs, HRP conjugated anti-human Fc polyclonal antibodies (Invitrogen, Thermo Fisher Scientific Inc., Waltham, MA) were added at 1:20,000 dilutions. Secondary antibodies were allowed to incubate for 1 hour at room temperature in the dark. After unbound secondary antibodies were washed thoroughly 5x with PBST, 3,3′,5,5′-Tetramethylbenzidine (TMB) substrate (R&D system, Minneapolis, MN) was added for 10 mins at room temperature to develop the chromogenic signal. Signal development was quenched with a Stop Solution (R&D system, Minneapolis, MN), and signal absorbance was measured at 450nm wavelengths with absorbance at 550nm as a background reference wavelength. Raw data was processed with Graphpad Prism, using a 4PL nonlinear regression fitting to calculate binding EC50 values.

### Competitive ELISA

Recombinant antigens of interest were immobilized onto ELISA plates as previously described. After washing with PBST, excess ligand was added to the plate with low concentrations of Nb or Nb-Fc fusion constructs (25nM and 1nM, respectively). 10μM transferrin was added for TfR1 competition experiments and 0.5μM HGF was used for cMET. Competitive binding was allowed to occur at room temperature for 2 hours. For non-Fc fused Nb constructs, HRP conjugated anti-T7 polyclonal antibodies (Invitrogen) were added at 1:7,500 dilutions. For Fc-fused Nb constructs, HRP conjugated anti-human Fc polyclonal antibodies (Invitrogen) were added at 1:20,000 dilutions. Secondary antibodies were allowed to incubate for 1 hour at room temperature in the dark. ELISA plates were washed and TMB substrate signal was developed as previously described. Antigen-coated wells with Nb and no ligand were used as 100% signal reference controls, and serial dilutions of Nb constructs were used to ensure that signals were within quantifiable linear range.

### Nb Affinity Measurement by SPR

Surface plasmon resonance (SPR, Biacore 3000 system, GE Healthcare, Chicago, IL) was used to measure Nb affinities. Nb was immobilized on the activated CM5 sensor-chip in pH 5.5 10mM sodium acetate buffer. The surface of the sensor was then blocked by 1 M ethanolamine-HCl (pH 8.5). A series of dilution of ECD of hTfR1 (spanning three orders of magnitude) was injected in HBS-EP+ running buffer (GE-Healthcare, Chicago, IL), at a flow rate of 20 μl/min for 180 s, followed by a dissociation time of 15 mins. Between each injection, the sensor chip surface was regenerated with 50mM NaOH. The regeneration was performed with a flow rate of 40 μl/min for 30 s, with 2.5 min equilibration time in buffer. The measurements were duplicated. Binding sensorgrams for Nb were processed and analyzed using BIA evaluation by fitting with the 1:1 Langmuir model or the 1:1 Langmuir model.

### Confocal Microscopy

Briefly, 1.5 x 10^5^ NCI-H1975 cells were plated in a NUNC^TM^ 96-well optical-bottom microplate (Thermo Fisher Scientific Inc., Waltham, MA) and treated with Tritazumab conjugated with Alexa Flour^TM^ 647 C2-maleimide (Thermo Fisher Scientific Inc., Waltham, MA). The plate was visualized using Leica TCS SP8 AOBS (Leica Microsystems, Germany). The objective lens specification is HC PL APO CS2 40x/1.30 oil immersion and an immersion oil F (RI=1.518) was used for imaging. The following specification were used for the imaging: (1) DAPI (blue), laser power: 2.00%, PMT detector spectral range: 410-567nm, PMT detector gain: 493.1V (2) Wheat germ (green), laser power: 0.25%, HyD detector spectral range: 567-628nm, HyD detector gain: 10.0% (3) 647 (red), laser power: 1.00%, HyD detector spectral range: 638-776nm, HyD detector gain: 58.0%.

### Flow Cytometry Cell Surface Binding

NCI-H1975 and HFL-1 cells were detached from growth dishes and resuspended at 100,000 cells/well in a 96-well V-bottom plate (Thermo Fisher Scientific Inc., Waltham, MA) with PBSA (1x PBS buffer with 0.1% Bovine 617 Serum Albumin). Cells were incubated for 3 hours with rotation at room temperature with titrations of Nb-Fc or antibody. Cells were washed with cold PBSA and incubated with goat anti-human Fc-Alexa Fluor 647 secondary antibody (Thermo Fisher Scientific Inc., Waltham, MA) or goat anti-human IgG H+L-Alexa Fluor 488 (Thermo Fisher Scientific Inc., Waltham, MA). After a PBSA wash, cells were analyzed for protein binding using an Attune flow cytometer. Data was analyzed using FlowJo (BD Biosciences, Franklin Lakes, NJ) and plotted using GraphPad Prism software. Binding curves were fitted to a logistic regression model. Experiments were performed in duplicates.

### Flow Cytometry Degradation Studies

NCI-H1975 and NCI-H1650 cells were grown in NUNC^TM^ 6 well plates (Thermo Fisher Scientific Inc., Waltham, MA) in complete media. For dose-response degradation studies, wells were treated with different concentrations of Tritazumab or controls for 24 hours before being washed and detached. For time course degradation studies, 10 nM Tritazumab or controls were added to different wells at 1.5, 3, 6, 12, and 24 hours prior to harvest before being washed and detached from the plate. For both study types, cells were washed in PBSA and transferred to V-bottom plates at a cell density of 100,000 cells per well. Cells were incubated for 3 hours with AlexFluor488-conjugated S11-Fc, 7D12-Fc, or Nb15-Fc in PBSA at room temperature for 3 hours with rotation, before a final PBSA wash and resuspension. Membrane protein expression levels were analyzed using an Attune flow cytometer. Data was analyzed using FlowJo (BD Biosciences, Franklin Lakes, NJ) and plotted using GraphPad Prism software. Experiments were performed in triplicates.

### Mouse Xenograft Studies

Female immunocompromised NU/NU B/C mice (10-12 weeks old) (Charles River, Wilmington, MA) were engrafted with 5 x 10^6^ NCI-H1650 cells per 200 μL in (PBS:matrigel=1:1). After tumor volumes reach ∼100 mm^3^, mice were randomized into different treatment arms and administrated with vehicle control or 1 mg/kg Tritazumab-MMAE at day 1 or day 12 via IP injections. The experimenters were not blinded. Tumor volume was calculated as follows: tumor size (mm^3^) = (longer measurement x shorter measurement^2^) x 0.5. Both tumor size and body weight were recorded every other day over the course of the studies. All procedures involving mice and experimental protocols (protocol ID: IACUC-2018-0013) were approved by the Institutional Animal Care and Use Committee (IACUC) of Icahn School of Medicine at Mount Sinai (ISMMS). Tumor growth inhibition was calculated using the following equation: [1 - (treatment)/(vehicle)] x 100. Blood was collected on day 21 for complete blood count (CBC) analysis was performed by the Center for Comparative Medicine and Surgery (CCMS) laboratory services at ISMMS. Data was analyzed using GraphPad Prism 8 and statistical analysis was performed using unpaired two-tailed *t*-tests.

Female NSG mice (8-10 weeks old) were engrafted with 5 x 10^5^ MDA-MB-231 cells suspended in 100 μL of PBS/Matrigel (9:1). After tumor volumes reached ∼30 mm^3^, mice were randomized into different treatment arms and administered with vehicle control or 1 mg/kg Tritazumab-ZZ7-16-073 on days 20, 23 and 27 via IP injections. The experimenters were not blinded. Tumor volume was calculated as follows: tumor size (mm^3^) = (longer measurement x shorter measurement^2^) x 0.5. Tumor volume and body weight were recorded every other day throughout the study. All procedures involving mice and experimental protocols (IACUC number: #23109) were approved by the Institutional Animal Care and Use Committee (IACUC) of City of Hope. Tumor growth inhibition was calculated using the following equation: [1 - (treatment)/(vehicle)] x 100. Data were analyzed using GraphPad Prism 8 and statistical analysis was performed using two-way ANOVA with multiple comparison test.

### Computational Molecular Interaction Study

We used the AlphaFold3 web server (https://alphafoldserver.com) to predict the complex structure of hTfR1 and S11.^31^ The amino acid sequence of hTfR1(uniprot: P02786, residues 122 to 760) and S11 were provided to AlphaFold3 web server. In the AlphaFold3 prediction, hTfR1 was modeled in its monomeric state.

### Statistical analysis

Statistical analyses were performed using GraphPad Prism 8 (GraphPad Software). A two-tailed Student’s t-test was used to compare the differences between two groups, represented as *P < 0.05, **P < 0.01 and ***P < 0.001; P > 0.05 was considered not significant.

## Reporting summary

Further information on research design is available the Nature Portfolio Reporting Summary linked to this article.

## Data availability

All data that support the findings of this study are available within the article and Supplementary Information or from the corresponding authors upon reasonable request.

## Code availability

The paper does not report original code.

## Author information

M.K, J.K., Z.D.,Y.Xiang, and P.R.S. contributed equally to this work.

## Author Contributions

Y.S., M.K., YJ.K., and J.J. conceived the study. Y.S., M.K., YJ.K. drafted the manuscript with substantial input from Y.Xiang, Y.Xiong, P.R.S, and J.J. J.J and Y.S. supervised the study. M.K. performed immunoblotting, cell viability, IHC studies. YJ.K. designed, cloned, and produced Tritazumab and its derivative antibody constructs, optimized and produced drug conjugates for *in vitro* and in vivo studies, and performed ELISA experiments. Y.Xiang, YJ.K, and Y. Xiong performed quality control experiments of antibody constructs (Q-TOF, SDS-PAGE, FPLC). Y.Xiang discovered and characterized cMET and TfR1 Nbs, and measured S11 affinity by SPR. Z.S. modeled S11-TfR1 interaction. C.J. initiated cMET camelid immunization and performed ELISA binding experiments. Z.D performed the chemistry synthesis. P.R.S. performed flow cytometry cell-surface binding and receptor degradation studies. R.H., and Y.L. performed related biochemical and cellular assays. A.B. purified Tritazumab for Q-TOF analysis with the help of Y. Xiang. M.K., N.J.P., A.Y., and Y.Z. designed and performed the in vivo study of Tritazumab-MMAE under the guidance of E.G., Z.W and K.H performed gene expression analysis. Y. Xiong supervised the chemistry synthesis and Q-TOF studies. N.S performed the in vivo study of Tritazumab-ZZ7-16-073 under the supervision of M.F.

## Acknowledgement

J.J. acknowledges the support by an endowed professorship by the Icahn School of Medicine at Mount Sinai (ISMMS). This work utilized the NMR Spectrometer Systems at Mount Sinai acquired with funding from NIH SIG Grants 1S10OD025132 and 1S10OD028504. M.K. acknowledges the support by the NIH-funded post-doctoral training grant in Cancer Biology (T32CA078207) at ISMMS. YJ.K acknowledge support by the NIH-funded pre-doctoral training grant in Cancer Biology (T32CA078207), and pre-doctoral training grant in Pharmacological Sciences (T32GM062754). Y.Z. acknowledges the support by the NIH-funded pre-doctoral training grant in Cancer Biology (T32CA078207), pre-doctoral training grant in Pharmacological Sciences (T32GM062754), and post-doctoral training grant in Cancer Biology (T32CA078207) at ISMMS. M.F. acknowledges the support by a NIH grant (R01CA258778). This work was partially supported by faculty startup funds from the University of Pittsburgh School of Medicine, a grant from The Michael J. Fox Foundation (co-funded by the ALZN Foundation. Y.S. is grateful to William E. Klunk and Arthur S. Levine (University of Pittsburgh) for their support and inspiration.

**Correspondence and requests for materials** should be addressed to Md Kabir, Jian Jin and Yi Shi.

