## Supplemental figures for "Multispecific nanobody degraders co-deplete membrane receptors and enable targeted delivery of diverse payloads"

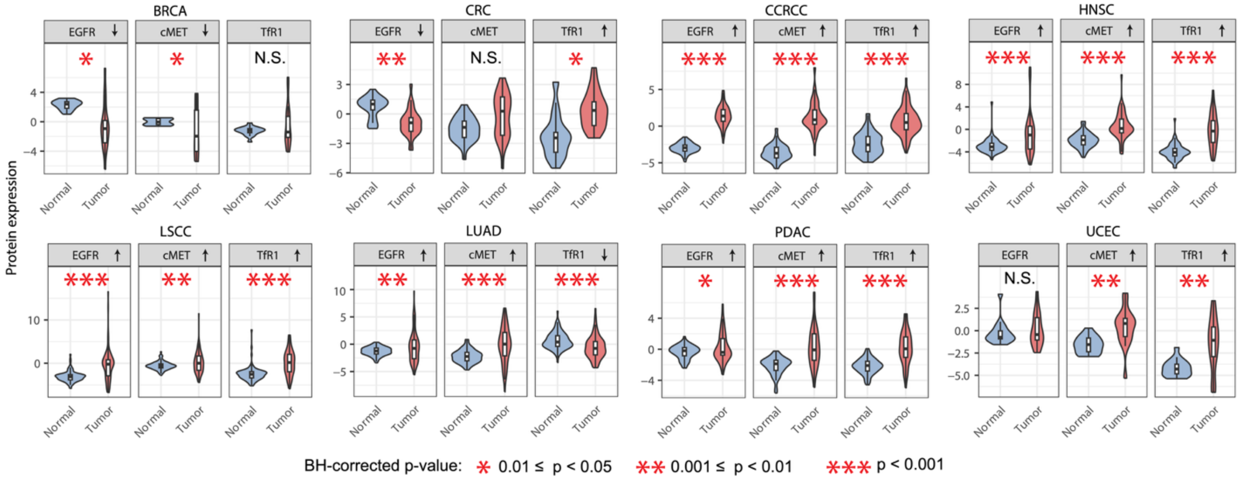


**Figure S1.** Overexpression of EGFR, cMET and TfR1 in solid cancers. CPTAC proteomics database was used to assess the protein expression of the EGFR, cMET and TfR in different cancer types compared to its normal tissue subtypes.

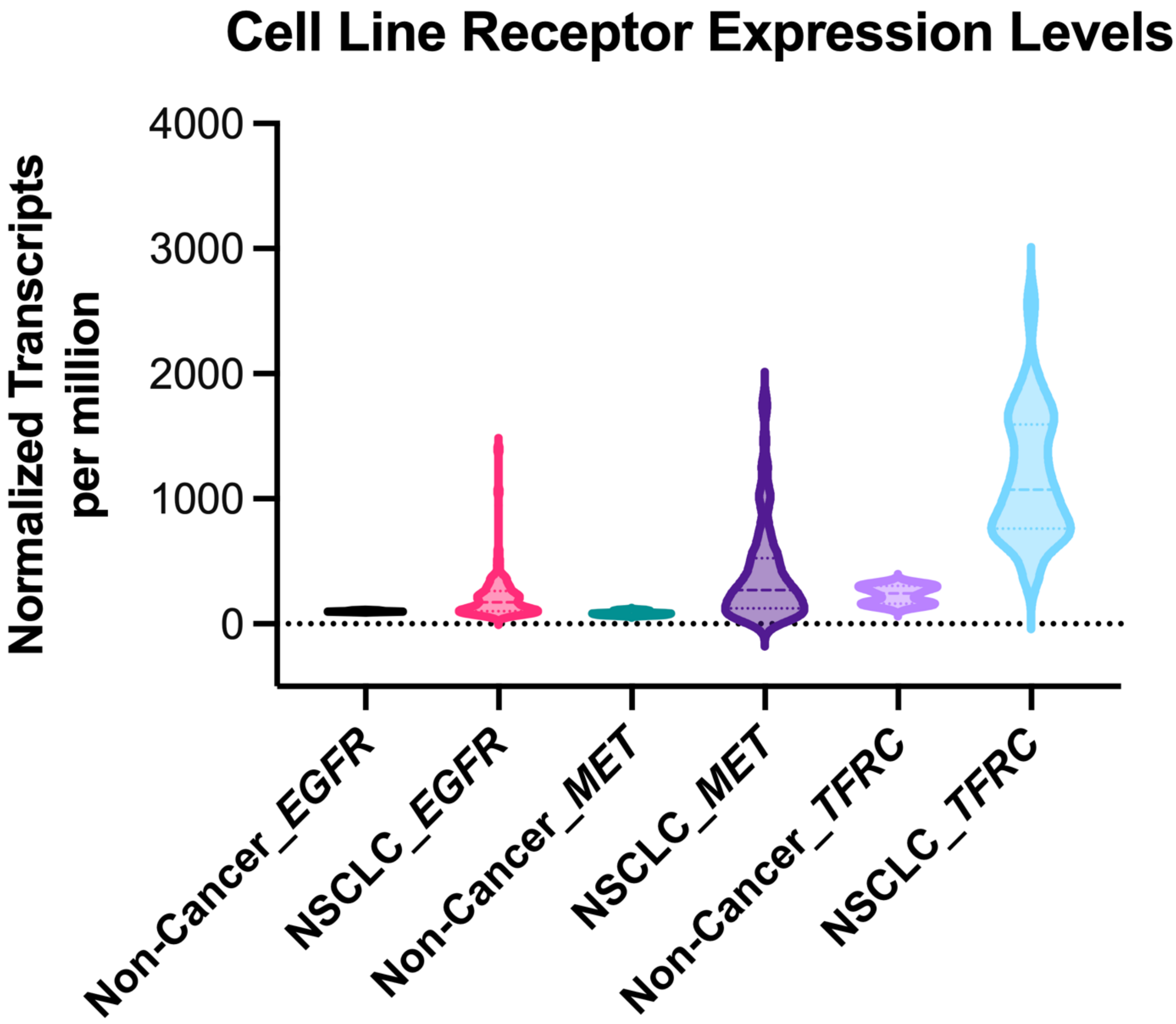


**Figure S2**. Gene expression of *EGFR*, *MET* and *TFRC* in NSCLC and non-cancer cell lines. MERAV database was utilized for the generation of the expression profile for *EGFR*, *MET* and *TFRC* receptors in NSCLC and non-cancer cell lines.

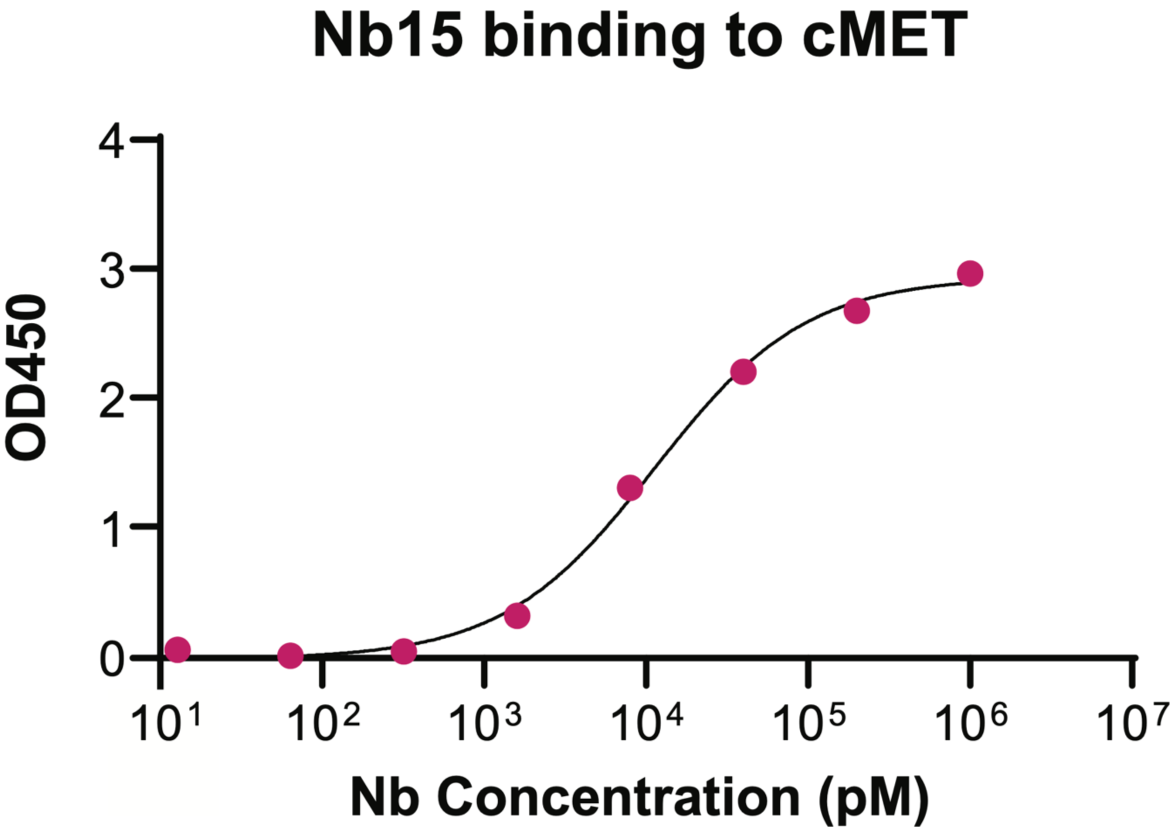


**Figure S3**. Binding assessment of a cMet-targeting Nb by ELISA (N = 1).

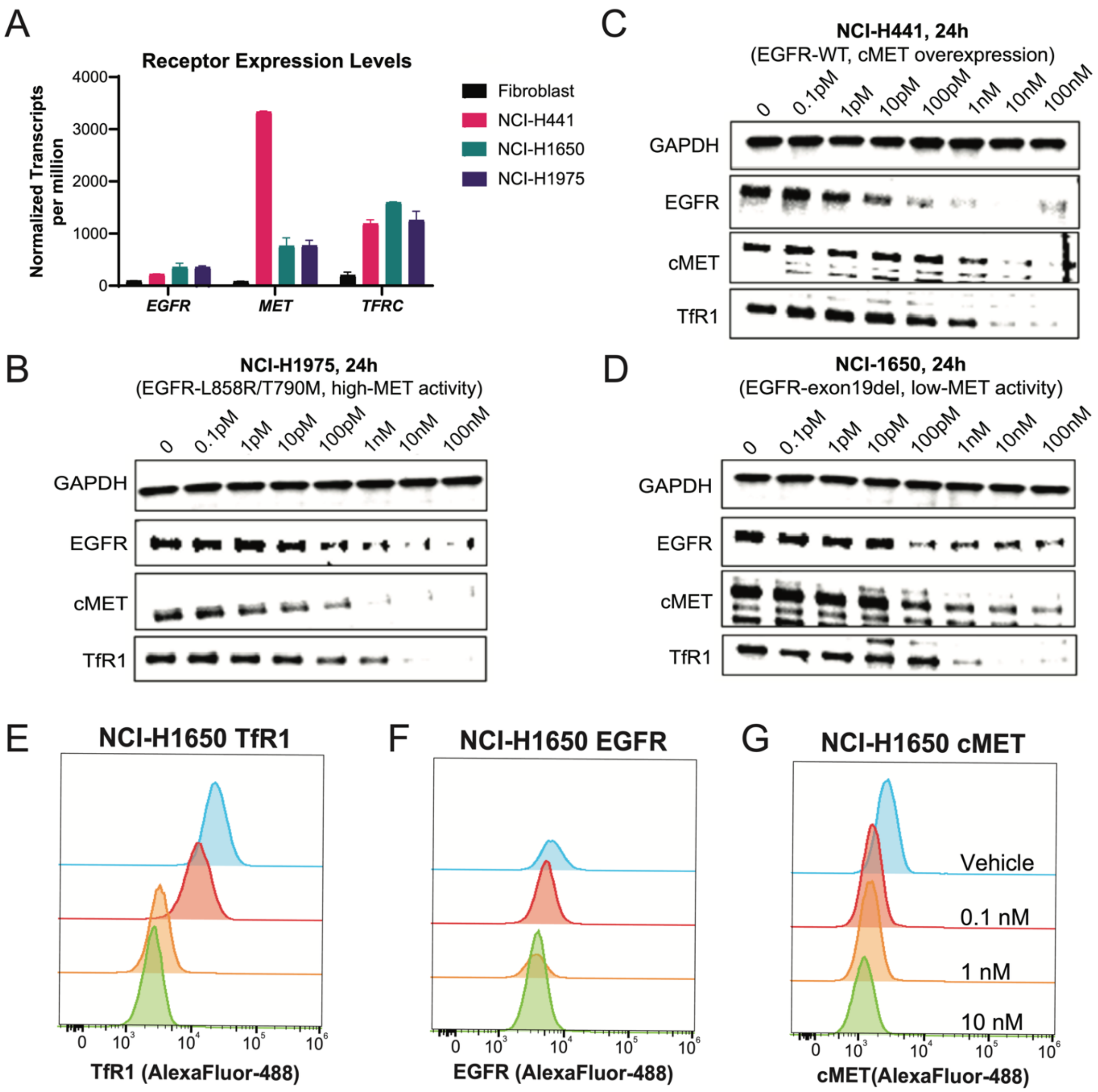


**Figure S4**. Characterization of Tritazumab in NSCLC cell lines. (A) *EGFR*, *MET* and *TFRC* receptor gene expression levels in lung fibroblast, NCI-H441, NCI-H1650 and NCI-H1975 cell lines. WB showing dose-dependent degradation of EGFR, cMET and TfR1 by Tritazumab in (B) NCI-H1975, (C) NCI-H441 and (D) NCI-H1650 cell lines after 24h treatment. (E-G) Flow cytometry assessment of cell-surface TfR1 (E), EGFR (F) and cMET (G) degradation by Tritazumab at 0.1, 1 and 10 nM concentration after 24h compared to vehicle (10 nM) in NCI-1650 cells and detected using AF488-conjugated Nb-Fc constructs.

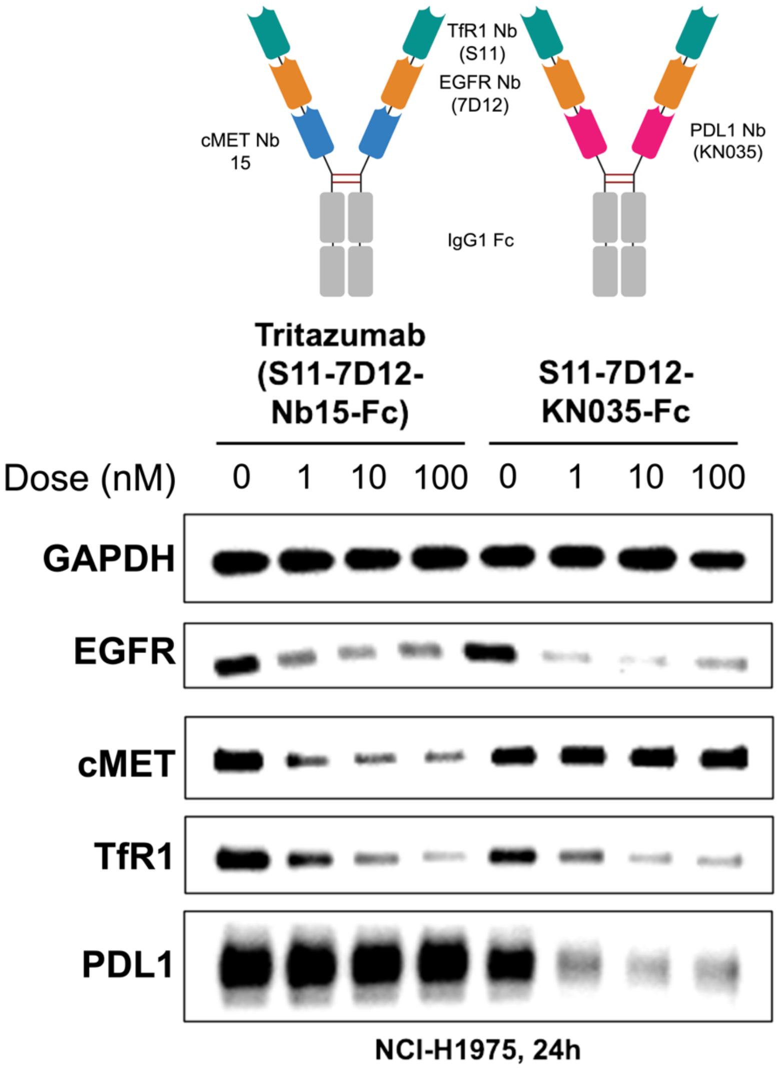


**Figure S5**. Development of a new MINDS construct (S11-7D12-KN035-Fc) targeting PD-L1, cMET and TfR1. WB analysis of EGFR, cMET, TfR1 and PD-L1 protein levels in NCI-H1975 cells treated by different MINDS constructs . Briefly, each construct was treated at 0, 1, 10 or 100nM in NCI-H1975 cell lines for 24h. Total cell lysates were used for WB and GAPDH was used as a loading control. The results are representative of 2 independent experiments.

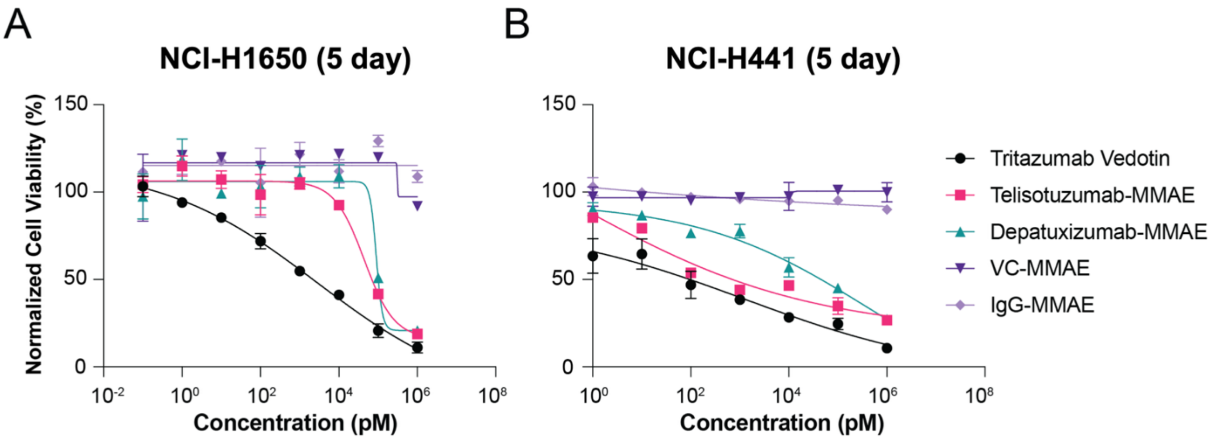


**Figure S6.** Cell viability of Tritazumab-MMAE, Telisotuzumab-vedotin, Depatuxizumab-MMAE, IgG-MMAE and VC-MMAE in (A) NCI-H1650 and (B) NCI-H441 cells. The cells were treated with vehicle or the indicated compounds at the indicated concentrations for 5 days. The mean $IC_{50}$ value ± SD for each concentration point (N = 3) is shown.

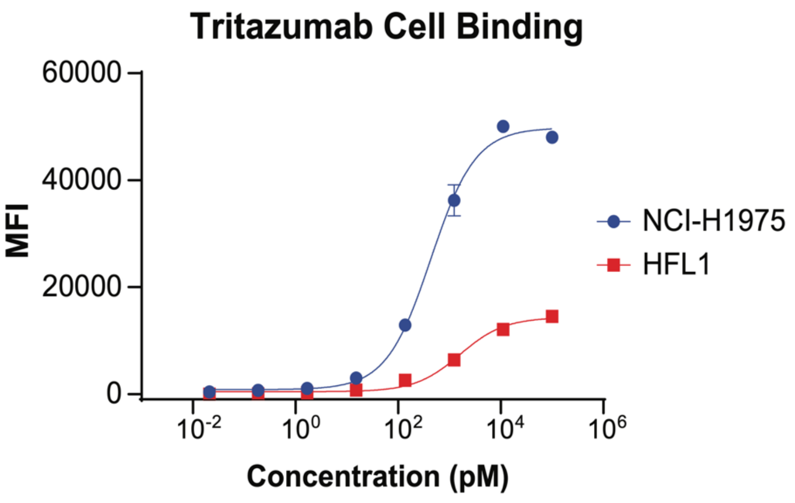


**Figure S7.** Tritazumab cell-binding comparison studies. Flow cytometry cell-binding of Tritazumab against cancerous NCI-H1975 cells and normal human lung fibroblast HFL1 cells (N=2).

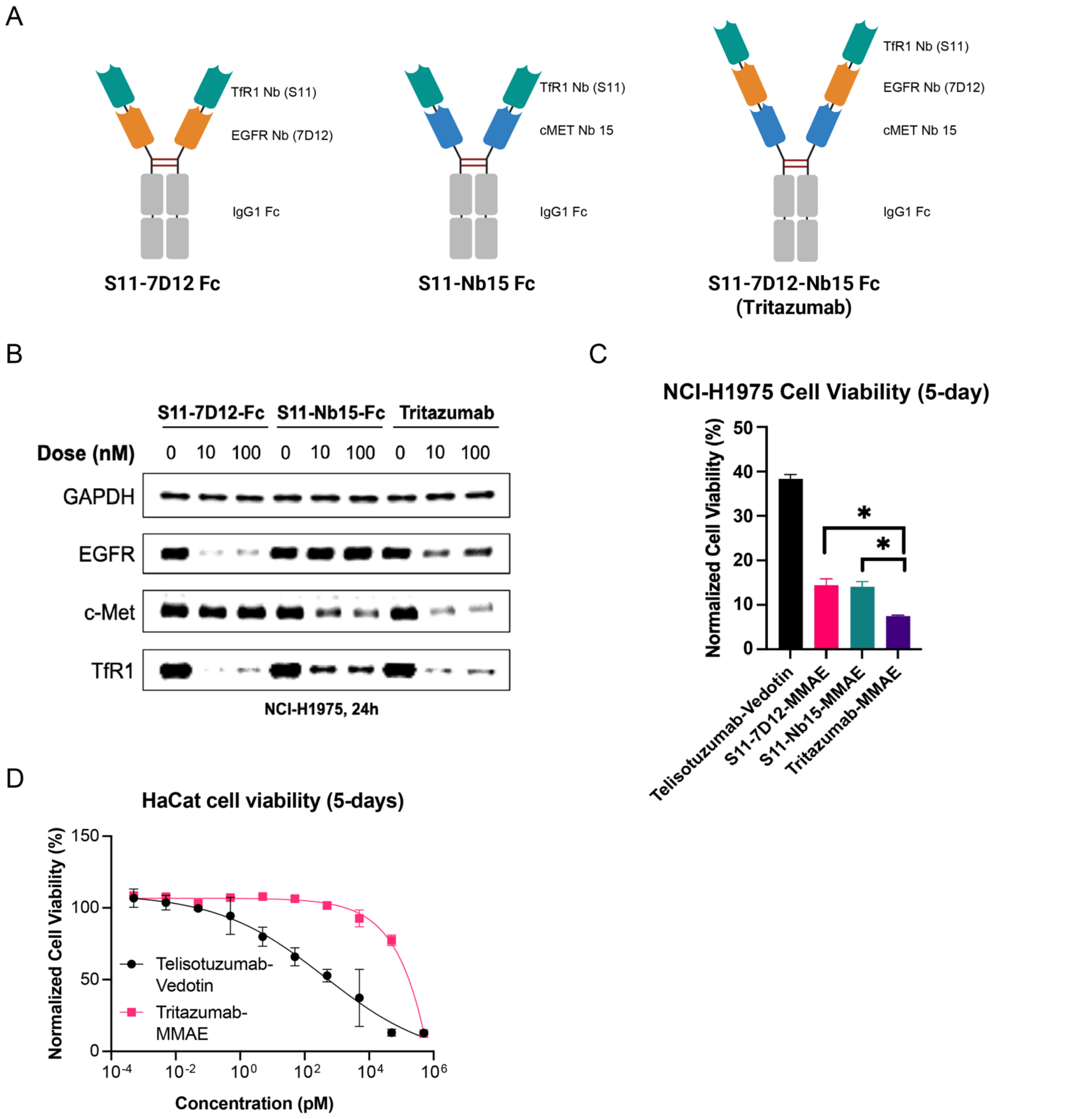


**Figure S8.** Comparison of Tritazumab to bispecific constructs and skin cell toxicity studies. (A) Schematic of S11-7D12-Fc and S11-Nb15-Fc dimer constucts alongside Tritazumab (B) WB analysis of EGFR, cMET and TfR1 degradation by the dimer constructs compared to Tritazumab. Each construct was treated at 0, 10 or 100 nM in NCI-H1975 cell lines for 24h. Total cell lysates were used for WB and GAPDH was used as a loading control. WB results shown are representative of 2 independent experiments. (C) Normalized cell viability of Telisotuzumab-vedotin, S11-7D12-MMAE, S11-Nb15-MMAE and Tritazumab-MMAE in NCI-H1975 cells at 1nM. The cells were treated with vehicle or the indicated compounds at the indicated concentrations for 5 days (N = 2 biological experiments). (D) Cell viability of Tritazumab-MMAE and Telisotuzumab Vedotin HaCat cells. The cells were treated with indicated compounds at the indicated concentrations for 5 days. The mean $IC_{50}$ value ± SD for each concentration point (N = 2).

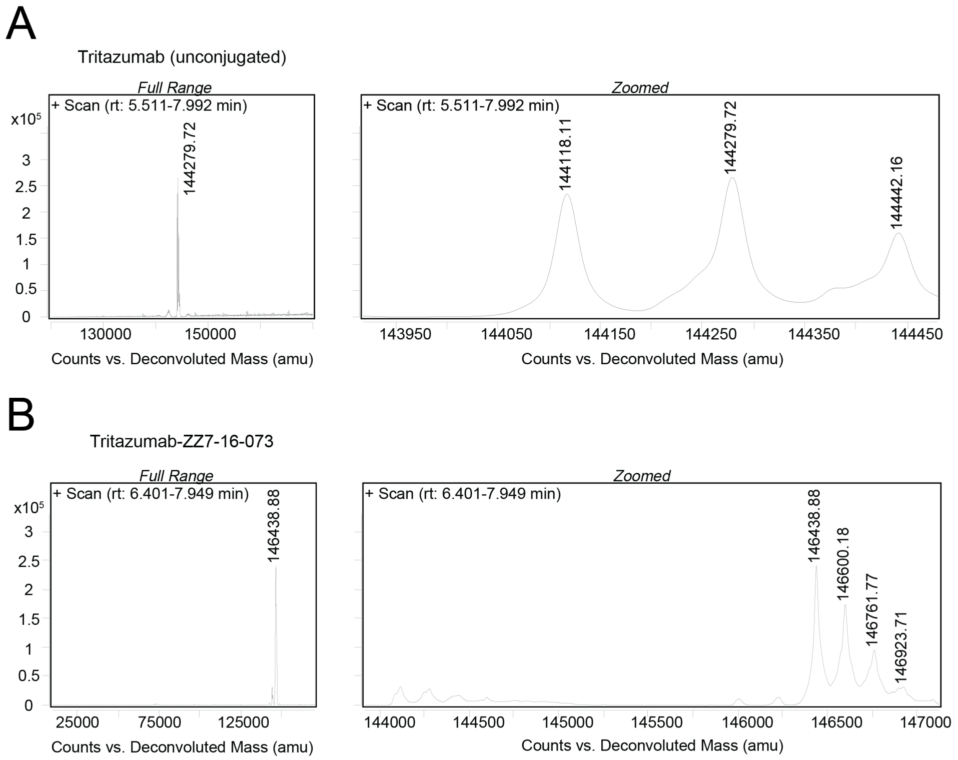


**Figure S9**. Q-TOF deconvoluted mass spectra of intact Tritazumab and Tritazumab–ZZ7-16-073. (A) Tritazumab exhibits a measured mass of 144,118.11 Da, consistent with G0F-type Fc N-glycosylation relative to the theoretical monoisotopic mass (141,400.48 Da). (B) Tritazumab–ZZ7-16-073 shows a mass of 146,438.88 Da, corresponding to a DAR of 2 (+2,318.96 Da, 12 ppm). Minor species spaced by ~162 Da are attributed to glycoform heterogeneity.

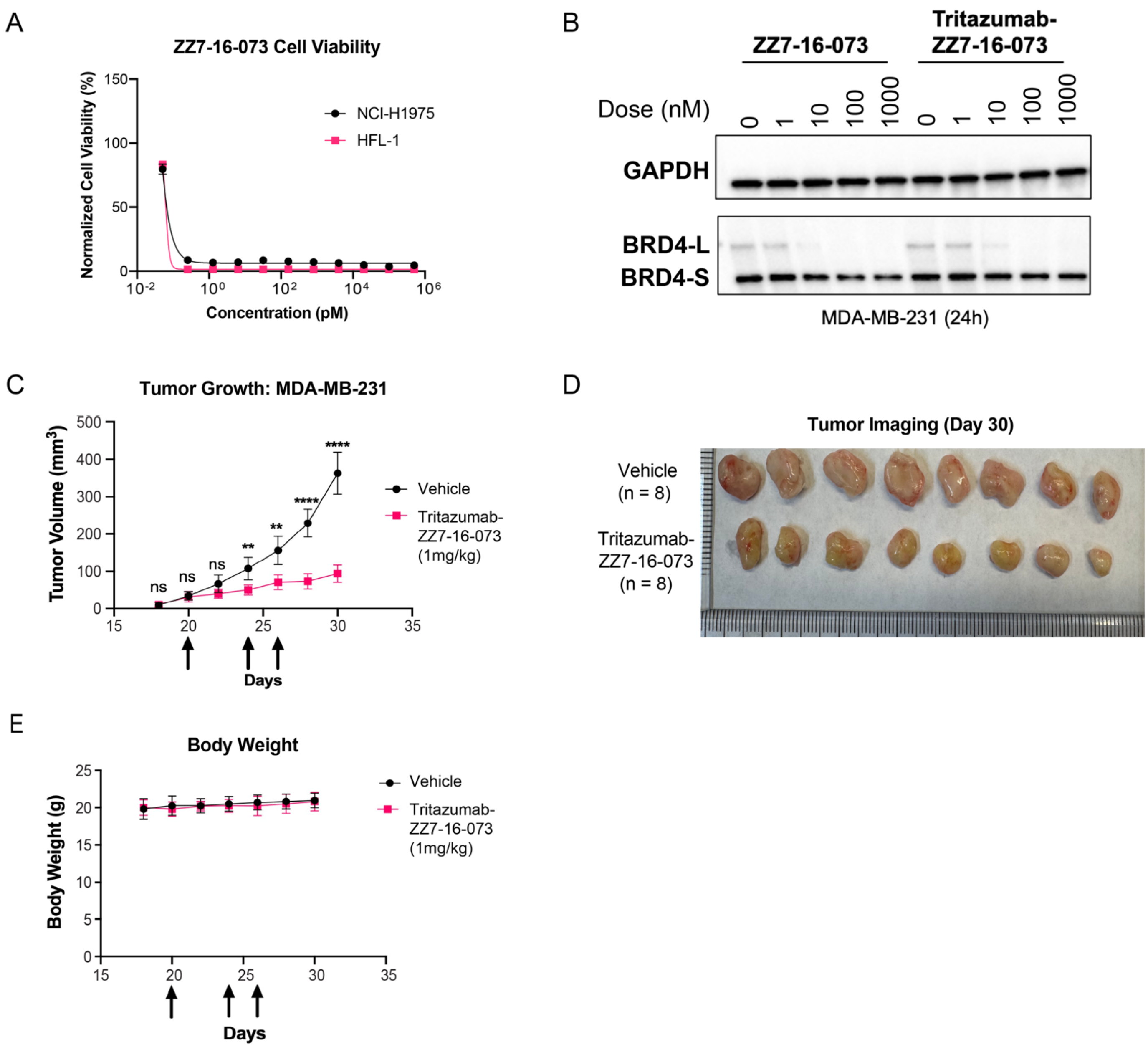


**Figure S10**. *In vitro* and *in vivo* evaluation of the Tritazumab-ZZ7-16-073 molecular glue conjugate. (A) Cell killing potency of ZZ7-16-073 in NCI-H1975 vs HFL-1 cell line. The cells were treated with vehicle or the indicated compounds at the indicated concentrations for 5 days. The mean value ± SD for each concentration point (in technical duplicates from three biological experiments) is shown in the curves. (B) WB results of BRD4 protein levels in MDA-MB-231 cells treated with ZZ7-16-073 or Tritazumab-ZZ7-16-073 at 0,1,10, 100 and 1000 pM for 24h (N = 2). Total cell lysate was used for WB and GAPDH was used a loading control. (C) Average tumor growth curves showing Tritazumab-ZZ7-16-073 led to significant tumor inhibition compared to vehicle groups. Data are presented as median values with standard errors of the mean from n = 8 mice for both treatment and control groups. P-values (ns - not significant, or *** - p-value<0.001) were determined by two-way ANOVA with multiple comparison test. (D) Images of harvested tumors on day 30. (E) Average weight measurement showing Tritazumab-ZZ7-16-073 does not lead to decreased body weight compared to vehicle groups.

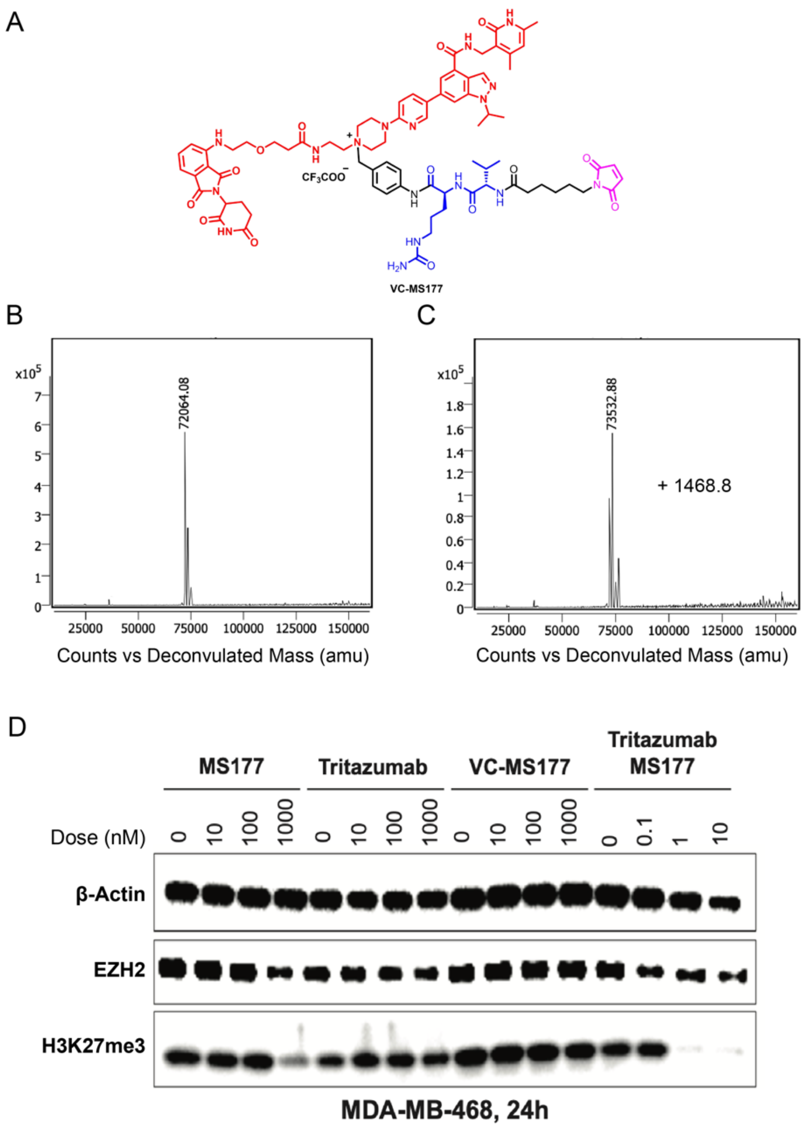

**Figure S11**. Development of Tritazumab-EZH2 PROTAC conjugate. (A) Structure of maleimide-containing EZH2 PROTAC, MS177 (termed VC-MS177). Q-TOF assessment of maleimide conjugation before (B) or after (C) VC-MS177 conjugation. (D) Evaluation of Tritazumab-MS177 conjugate compared to parental MS177, Tritazumab or VC-MS177 at 0, 10, 100 and 1000 nM treatment in MDA-MB-468 cell line for 24h (N = 2). Total cell lysate was used for WB and β-Actin was used a loading control.

### Supplementrary Experimental Section

**Chemistry General Procedures.** All commercially available chemical reagents were used directly in syntheses without further purification. A Teledyne ISCO Combi*Flash* Rf$^+$ instrument and HP C18 Redi*Sep* Rf reverse phase columns equipped with UV detector were used to conduct flash chromatography. All final compounds for biological evaluation were purified with preparative high-performance liquid chromatography (HPLC) on an Agilent Prep 1200 series with the UV detector set to 254 or 220 nm with a flow rate of 40 mL/min at room temperature. Crude samples were injected into a Phenomenex Luna 750 x 30 mm, 5 µ C18 column, with the gradient program set to 10% of methanol or acetonitrile (B) in $H_2O$ containing 0.1% TFA (A) progressing from 10 % to 100% of methanol or acetonitrile (B). The purity of all compounds for biological testing is > 95% assessed by HPLC-HRMS. All HPLC spectra was obtained by using an Agilent 1200 series system with DAD detector and a 2.1 mm x 150 mm Zorbax 300SB-C18 5 µ column for chromatography. Samples (0.5 µL) were injected onto a C18 column at room temperature with the flow rate of 0.4 mL/min. Chromatography was performed with the solvent as follows: water containing 0.1% formic acid was designated as Solvent A while acetonitrile containing 0.1% formic acid was designated as solvent B. The linear gradient was set such that 1% B was used from 0-1 min, 1-99% B from 1-4 min, and 99% B from 4-8 min. High-resolution mass spectra (HRMS) data was acquired in positive ion mode using an Agilent G1969A API-TOF with an electrospray ionization (ESI) source. All compounds were also characterized using either a Bruker (Billerica, MA) DRX Nuclear Magnetic Resonance (NMR) spectrometer (400 MHz, $^1$H NMR, 101 MHz $^{13}$C NMR). Chemical shifts for all compounds are reported in units of parts per million (ppm, δ) relative to residual solvent peaks. $^1$H NMR data are reported in the following format: chemical shift, multiplicity (s = singlet, d = doublet, t = triplet, q = quartet, m = multiplet), coupling constant, and integration. **Linker 1 – 12,**[1] and **I–2**[2] were prepared following the reported procedures.

**Synthesis of (*S*)-6-(2-(*tert*-butoxy)-2-oxoethyl)-8-(4-((*S*)-2-((*S*)-2-(6-(2,5-dioxo-2,5-dihydro-1*H*-pyrrol-1-yl)hexanamido)-3-methylbutanamido)-5-ureidopent**

**anamido)benzyl)-4-(4-(2-fluoro-5-nitrobenzamido)phenyl)-2,3,9-trimethyl-6*H*-thieno[3,2-*f*][1,2,4]triazolo[4,3-*a*][1,4]diazepin-8-ium (VC-ZZ7-16-073)**

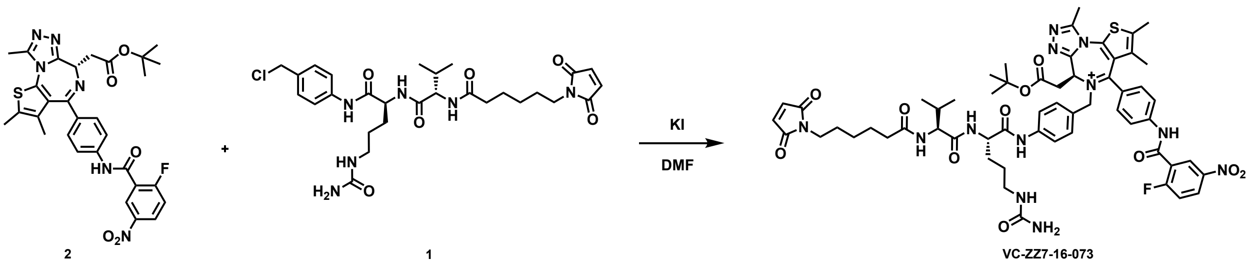

To a solution of compound **2** (12.2 mg, 0.02 mmol, 1.0 equiv)[3] and compound **1** (23.6 mg, 0.04 mmol, 2 equiv)[4] in DMF (0.2 mL) were added Potassium iodide(0.4mg, 0.002 mmol, 0.1 equiv). The reaction was stirred at 60 °C for 24h, then the resulting mixture was purified by preparative HPLC (10%-100% acetonitrile / 0.1% TFA in $H_2O$) to give the product **VC-ZZ7-16-073** (solid, 7 mg, 30%). $^1$H NMR (400 MHz, Methanol-*d*$_4$) δ 8.66 – 8.57 (m, 1H), 8.52 – 8.41 (m, 1H), 8.26 (d, *J* = 7.4 Hz, 1H), 7.77 (d, *J* = 8.6 Hz, 2H), 7.70 – 7.64 (m, 2H), 7.52 (d, *J* = 8.9 Hz, 3H), 7.45 (d, *J* = 8.4 Hz, 2H), 6.73 (s, 2H), 5.59 (d, *J* = 5.1 Hz, 2H), 4.71 (t, *J* = 7.2 Hz, 1H), 4.51 – 4.39 (m, 1H), 4.14 – 4.05 (m, 1H), 3.48 – 3.35 (m, 4H), 3.24 – 3.03 (m, 5H), 2.53 (s, 3H), 2.27 (t, *J* = 7.3 Hz, 2H), 2.13 – 1.99 (m, 1H), 1.92 – 1.83 (m, 1H), 1.80 (s, 3H), 1.77 – 1.68 (m, 1H), 1.63 (dd, *J* = 20.3, 7.3 Hz, 4H), 1.55 – 1.49 (m, 2H), 1.47 (s, 9H), 1.36 – 1.18 (m, 2H), 0.96 (d, *J* = 6.8 Hz, 6H). $^{13}$C NMR (101 MHz, Methanol-*d*$_4$) δ 176.56, 174.24, 172.69, 172.63, 171.32, 166.96, 165.66, 163.08, 162.43, 156.08, 153.05, 145.71, 142.43, 140.85, 136.95, 135.39, 134.74, 133.04, 131.11, 130.87, 129.59, 129.11, 127.41, 127.36, 126.67, 126.49, 121.49, 121.13, 119.27, 119.02, 82.93, 60.82, 55.71, 55.19, 54.35, 38.52, 38.23, 36.58, 31.83, 30.43, 29.48, 28.61, 28.01, 27.59, 26.53, 20.02, 19.15, 14.63, 13.40, 11.72. HRMS (ESI) m/z: calcd for $C_{58}H_{68}FN_{12}O_{11}S^+$ [M]$^+$, 1159.4830; found,1159.4854.

**Synthesis of 4-(5-(4-(((4,6-dimethyl-2-oxo-1,2-dihydropyridin-3-yl)methyl)carbamoyl)-1-isopropyl-1*H*-indazol-6-yl)pyridin-2-yl)-1-(4-((*S*)-2-((*S*)-2-(6-(2,5-dioxo-2,5-dihydro-1*H*-pyrrol-1-yl)hexanamido)-3-methylbutanamido)-5-ureidopentanamido)benzyl)-1-(2-(3-(2-((2-(2,6-dioxopiperidin-3-yl)-1,3-dio xoisoindolin-4-yl)amino)ethoxy)propanamido)ethyl)piperazin-1-ium (VC-MS177)**

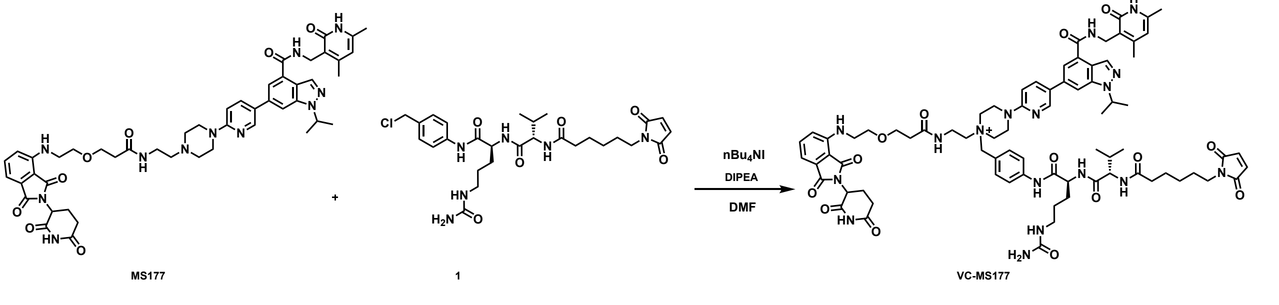

To a solution of **MS177** (26 mg, 0.025 mmol, 1.0 equiv) and compound **1** (16 mg, 0.0275 mmol, 1.1 equiv)[4] in DMF (0.2 mL) were added tetrabutylammonium iodide(4.7mg, 0.0125 mmol, 0.5 equiv) and DIEA (33uL, 0.188 mmol, 7.5 equiv). The reaction was stirred at rt for 24h, then the resulting mixture was purified by preparative HPLC (10%-100% acetonitrile / 0.1% TFA in $H_2O$) to give the product **VC-MS177** (solid, 7.4 mg, 20%). $^1H$ NMR (400 MHz, Methanol-$d_4$) δ 8.53 (s, 1H), 8.37 (s, 1H), 8.00 (d, $J$ = 8.1 Hz, 1H), 7.91 (s, 1H), 7.81 – 7.65 (m, 3H), 7.57 – 7.49 (m, 2H), 7.45 – 7.34 (m, 1H), 6.99 – 6.86 (m, 3H), 6.78 (s, 2H), 6.16 (s, 1H), 5.10 (s, 1H), 5.03 – 4.96 (m, 1H), 4.59 (t, $J$ = 37.3 Hz, 5H), 4.26 – 4.06 (m, 3H), 3.91 – 3.35 (m, 18H), 3.25 – 3.04 (m, 2H), 2.85 – 2.49 (m, 4H), 2.45 (s, 3H), 2.34 – 2.20 (m, 5H), 2.14 – 1.99 (m, 2H), 1.97 – 1.71 (m, 2H), 1.71 – 1.46 (m, 12H), 1.37 – 1.25 (m, 3H), 0.98 (s, 6H). $^{13}C$ NMR (101 MHz, Methanol-$d_4$) δ 176.64, 174.96, 174.77, 174.42, 172.95, 172.79, 172.19, 171.01, 169.30, 169.25, 165.83, 162.55, 153.89, 148.14, 145.22, 142.41, 141.47, 137.50, 137.05, 135.51, 135.31, 134.11, 133.66, 129.59, 128.53, 122.94, 122.74, 122.05, 121.45, 120.92, 120.47, 118.60, 117.64, 114.82, 112.42, 111.58, 111.07, 110.66, 70.66, 68.14, 65.94, 61.03, 58.35, 55.99, 55.31, 51.55, 50.37, 43.30, 40.56, 38.60, 38.36, 37.87, 37.18, 36.66, 34.24, 32.40, 31.77, 30.39, 29.52, 28.20, 27.59, 26.59, 23.92, 22.73, 20.03, 19.22, 18.84, 17.74. HRMS (ESI) m/z: calcd for $C_{76}H_{94}N_{17}O_{14}^+$ [M]$^+$, 1468.7161; found,1468.7137.

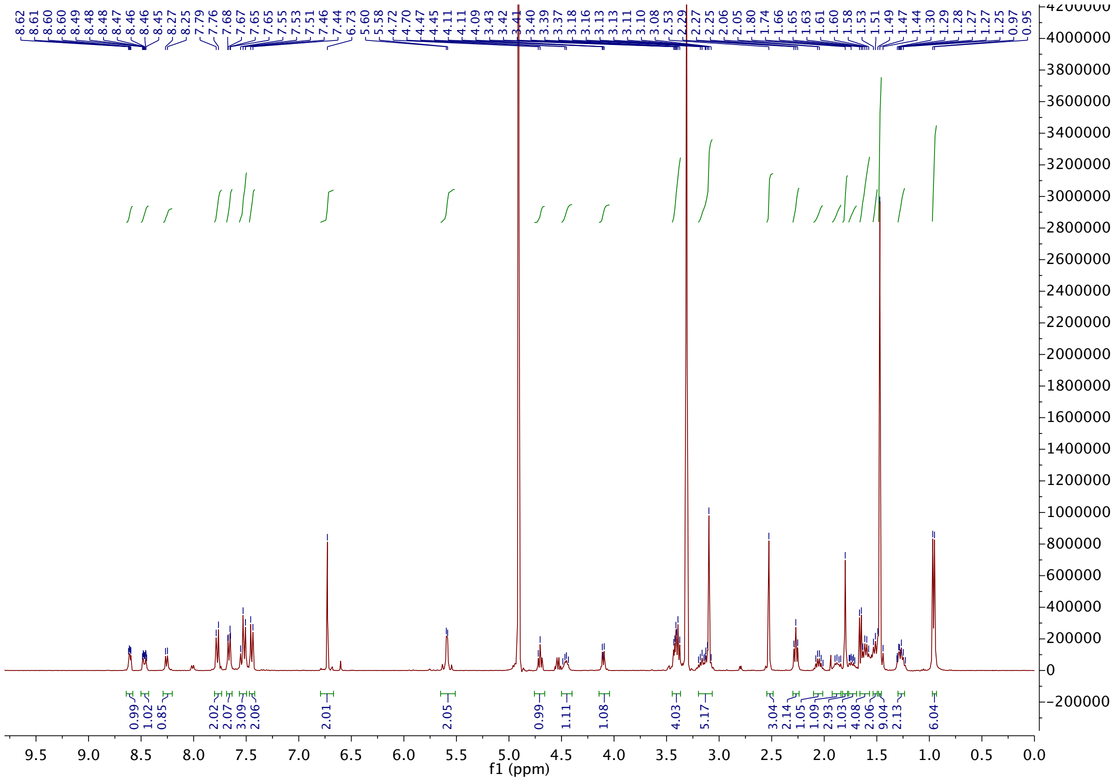

[1]H NMR spectrum of VC-ZZ7-16-073

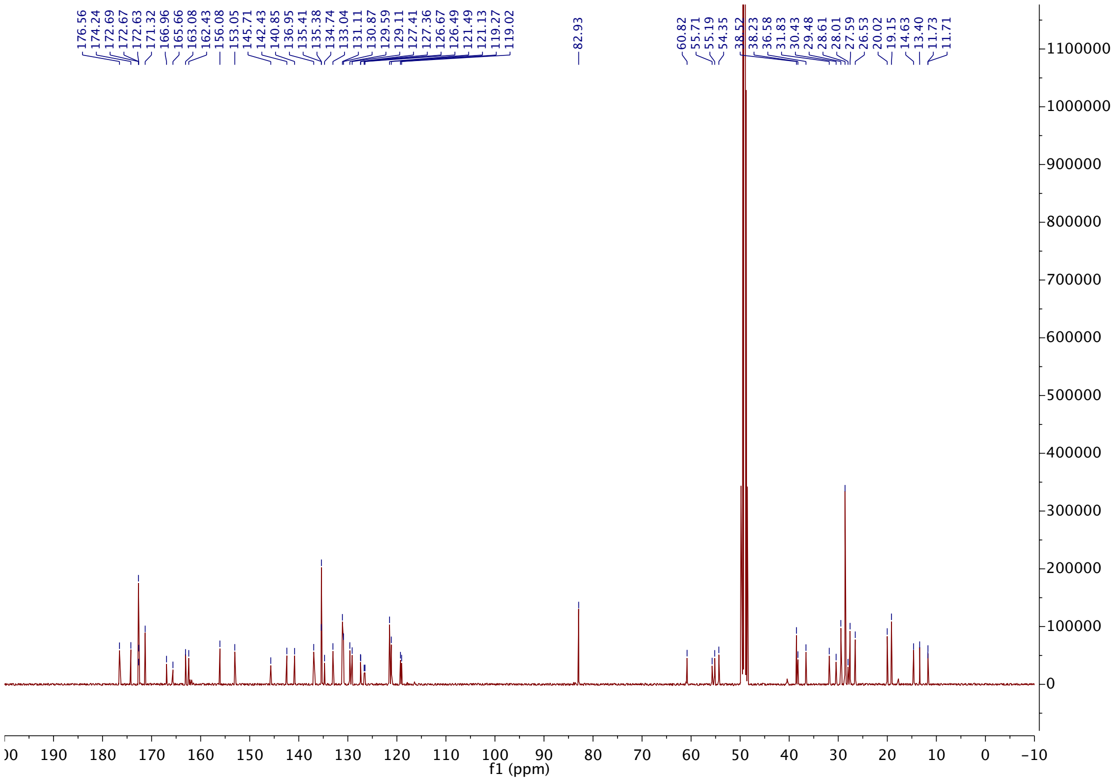


$^{13}C$ NMR spectrum of VC-ZZ7-16-073

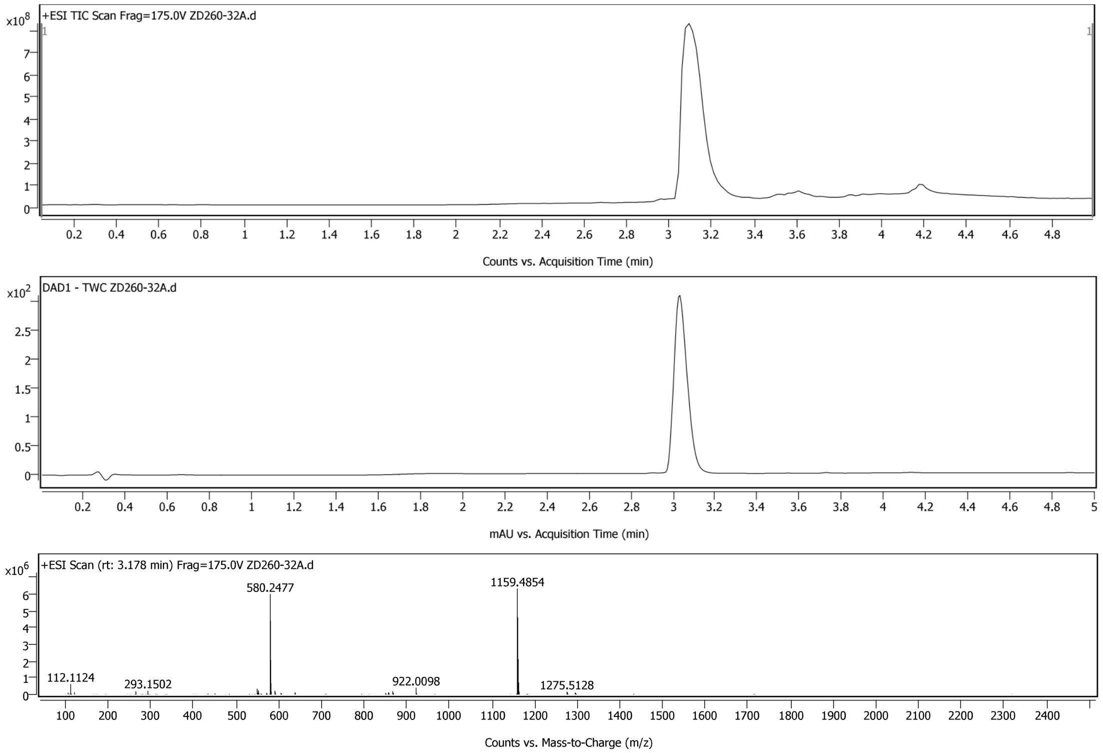


LC-MS spectra of VC-ZZ7-16-073

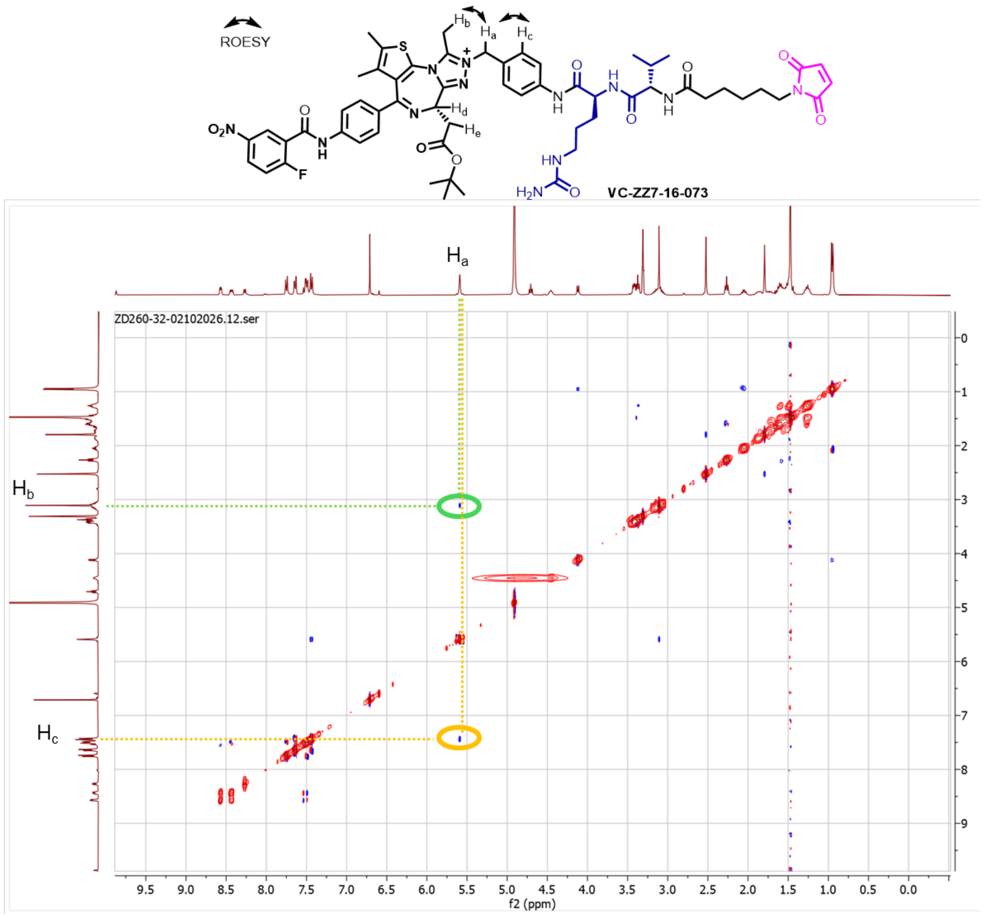


2D NMR ($^1$H-$^1$H ROESY) spectrum of VC-ZZ7-16-073

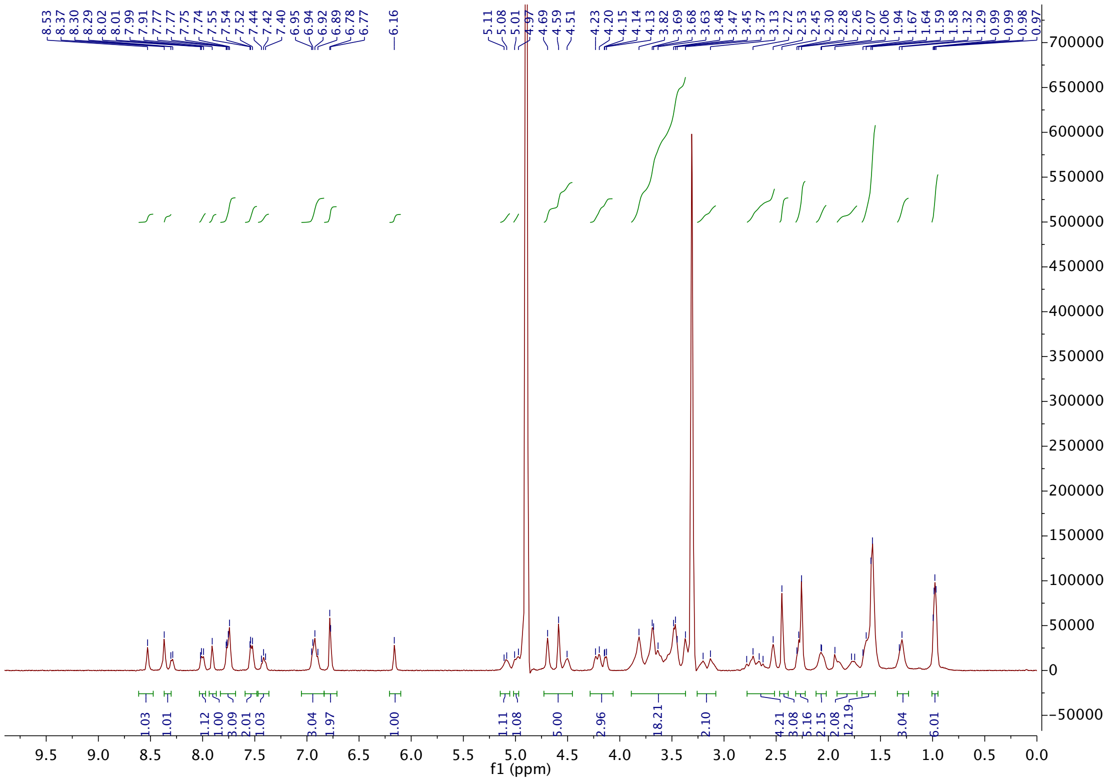

$^{1}$H NMR spectrum of VC-MS177

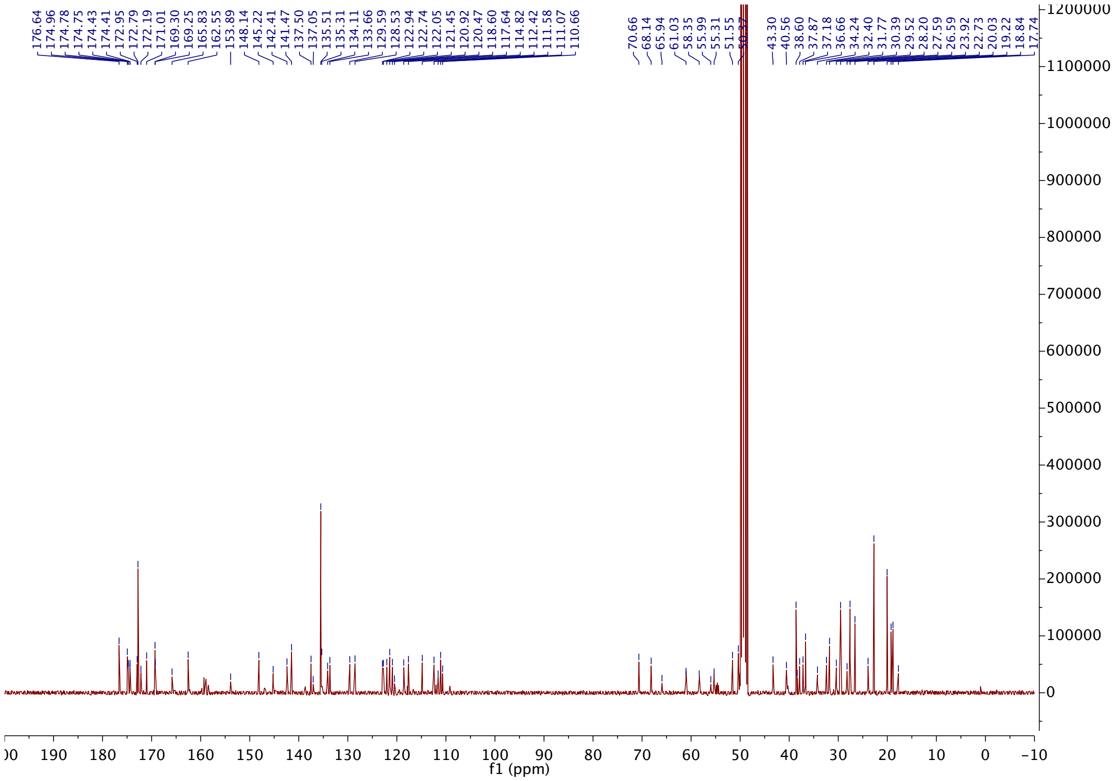

[13]C NMR spectrum of VC-MS177

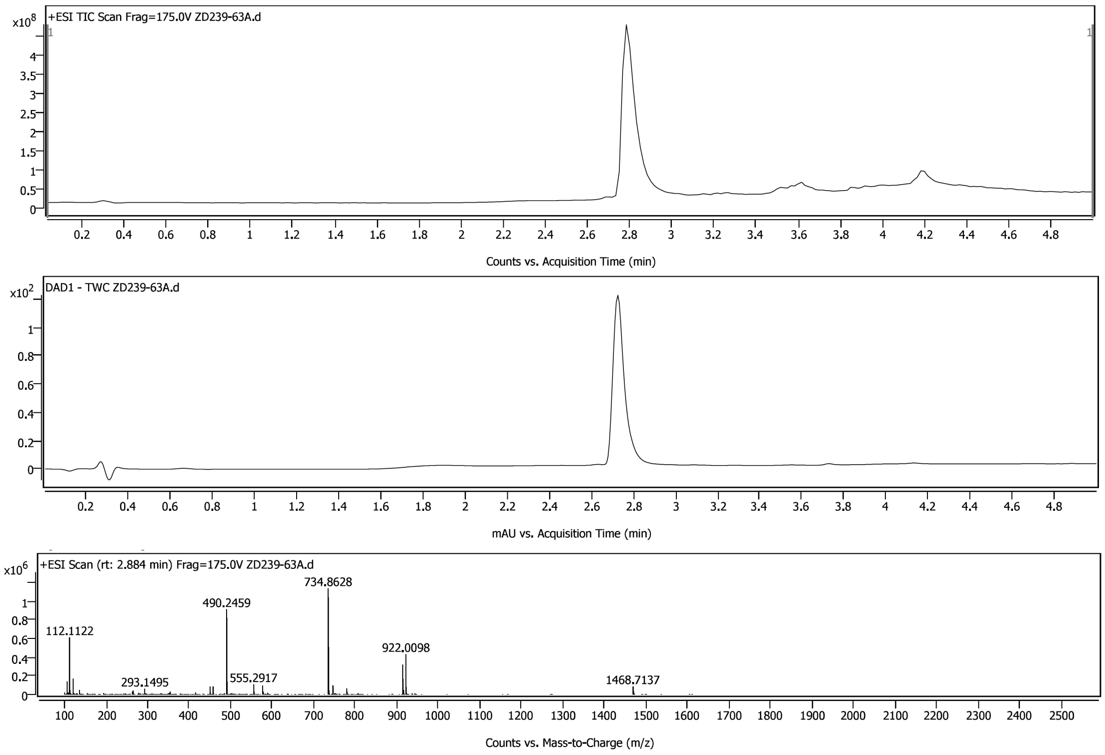


LC-MS spectra of VC-MS177

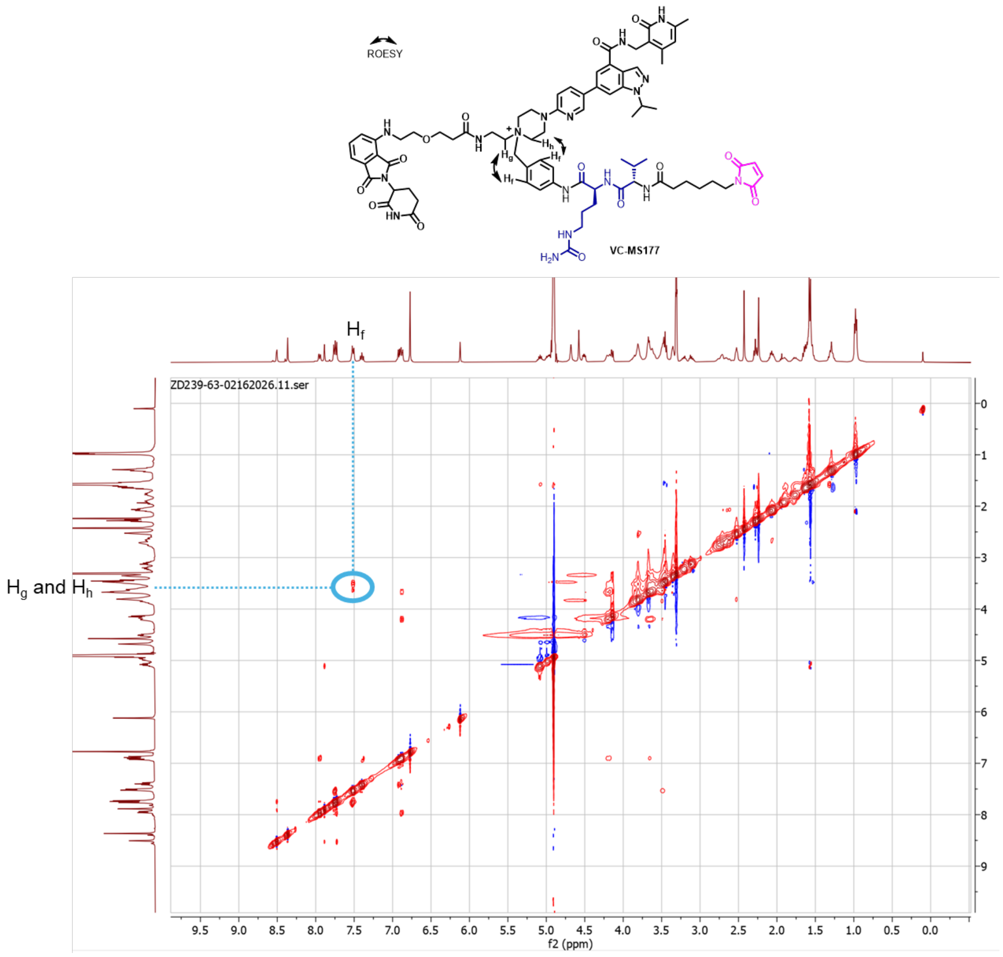


2D NMR ($^1$H-$^1$H ROESY) spectrum of VC-MS177
